# Comparative Genomic and Transcriptomic Analysis of Avirulent and Virulent Strains of *Fusarium oxysporum* f. sp. *carthami*: Insights into Pathogenesis and Virulence Determinants in Safflower Infections

**DOI:** 10.1101/2023.08.14.553163

**Authors:** Aabha, Vijay Laxmi, Babita Singh, Samriddhi Mehta, Ashish Kumar Gupta, Alok Srivastava, Samir Sawant, Surekha Katiyar-Agarwal, Kumar Paritosh, Manu Agarwal

**Affiliations:** Department of Botany, University of Delhi, Delhi, India; Centre for Genetic Manipulation of Crop Plants, University of Delhi, South Campus, Delhi, India; ICAR-National Institute for Plant Biotechnology, Pusa, New Delhi, India; CSIR-National Botanical Research Institute, Rana Pratap Marg, Lucknow, India; ICAR National Bureau of Agriculturally Important Microorganisms (NBAIM), Mau Nath Bhanjan, Uttar Pradesh, India; Department of Plant Molecular Biology, University of Delhi, New Delhi, India

## Abstract

Vascular wilt disease incited by *Fusarium oxysporum* f. sp. *carthami* (Foc) in Safflower poses a significant threat to its production in India. A comprehensive understanding of the molecular underpinning of compatible and incompatible interaction is of extreme economic importance. In the present study, the genome of a virulent (IARI-5175) and a avirulent (F-00845) Foc strain were sequenced and assembled using data generated from Illumina in combination with Nanopore technologies and HiC. Foc genomes were assembled into 88 and 23 scaffolds with an estimated total size of 46 Mb and 42 Mb respectively for IARI-5175 and F-00845 strains. Reference based mapping of Foc genome with *F. oxysporum* f. sp. *lycopersici* (Fol) resulted in chromosomal level ordering of genome and simultaneous identification of accessory genome. Additionally, two lineage specific chromosomes were also identified for virulent Foc strain IARI-5175. Genomic comparisons were made on the basis of effectors, CAZymes, secondary metabolites and mycotoxins to understand the global view pathogenicity in Foc. Moreover, the transcriptome of Foc during compatible and incompatible interaction was sequenced and analyzed leading to the identification of differentially regulated genes. Taken together our study laid a solid foundation to explore novel effector genes that play a crucial role in the establishment of disease and can further be used as targets to devise new strategies to curb wilt disease in safflower.

## Introduction

Safflower (*Carthamus tinctorius* L.), belonging to the Asterales: Asteraceae order, is a thistle-like, highly branched, herbaceous annual plant (Knowles, 1980). Traditionally grown for its seeds, vibrant flowers, and foliage, the seed oil of safflower is rich in oleic and linoleic fatty acids (mono- and polyunsaturated fatty acids respectively), making it a beneficial ingredient for heart health (Lee et al., 2004). Additionally, its flowers are often used for extracting natural dyes (Adamska & Biernacka, 2021). In India, safflower, known for its nutritious seed oil, is primarily cultivated as a rabi crop. The major safflower-producing states are Maharashtra, Karnataka, Andhra Pradesh and Telangana (Singh et al., 2019a). Despite India’s significant contribution to global safflower production, the crop is underutilized and remains a minor oilseed crop. Farmers are increasingly abandoning safflower cultivation in favor of alternative crops, leading to a continuous decline in annual safflower production. A primary reason for reduced safflower production in India is its susceptibility to fungal pathogens, notably *Alternaria carthami* and *Fusarium oxysporum* f. sp. carthami (Foc), the latter of which causes vascular wilt disease in safflower. The Fusarium wilt disease, instigated by Foc, poses a significant threat to safflower cultivation, reducing its yield by 40-80% annually (Singh et al., 2019b) and inflicts a considerable economic impact on Indian farmers. Foc initiates the wilt disease by infiltrating the plant’s roots, subsequently commandeering and blocking the vascular tissue xylem, resulting in severe wilting and plant death (Gordon, 2017).

*F. oxysporum*, a ubiquitous soil fungus classified within the Sordariomycetes, Hypocreales, and Nectriaceae families, may exist in a benign state as an endophyte (Nel et al., 2006; Wang et al., 2020a; J. Zhang et al., 2018) or metamorphose into a destructive pathogen, inflicting substantial damage to various crops of agricultural significance (Bishop & Cooper, 1983; Rishbeth, 1955). Even though *F. oxysporum* demonstrates an impressive adaptability to a wide array of ecological environments, many of its sub-species exhibit a high level of host-specificity and are accordingly classified into formae speciales (Edel-Hermann & Lecomte, 2019). Each formae speciales can include multiple strains, further categorized into pathogenic races predicated on their virulence potentials. The virulence of *F. oxysporum*, and its interaction with specific hosts, hinges on various elements like pathogenesis-related proteins, effector proteins, constituents of the signal transduction pathway, and transcription factors (TFs). Interestingly, numerous studies suggest that horizontal gene transfer allows *F. oxysporum* to modify its virulence characteristics and host specificity. This transfer of additional genomic regions enables *F. oxysporum* strains to rapidly evolve their virulence traits, all the while preserving essential genes vital for cellular processes. Thus, the variable evolution rates across separate genomic regions substantially influence the diversity noted in host range (including plants, animals, and immunocompromised humans) and host-specific pathogenicity (Fokkens et al., 2018; Frantzeskakis et al., 2019).

The genomes of *F. oxysporum* strains are structured into highly conserved core chromosomes and lineage-specific accessory chromosomes, the latter of which are conditionally essential for vegetative growth (Ma et al., 2010; Vlaardingerbroek et al., 2016). Despite the relative scarcity of genes and the prevalence of transposable elements and repeat-rich genomic regions within accessory chromosomes, they host several functional genes that endow the pathogen with an enhanced ability for virulence or host specificity. These chromosomes house genes associated with diverse biological processes, including metabolism, stress response, and signal transduction (Yang et al., 2020). Remarkably, these accessory chromosomes exhibit considerable plasticity, as they can be readily lost, gained, or transferred amongst strains during sexual or asexual reproduction. This flexibility appears to significantly contribute to the rapid evolution and adaptability of these fungal pathogens.

The location of effector genes on accessory chromosomes and their horizontal gene transfer between formae speciales serve as primary drivers in the pathogenomic evolution and adaptation of Fusarium strains to their hosts. Given the existence of diverse accessory genomes, it becomes imperative to sequence the entire genomes of strains from various races to thoroughly understand the complexity of genes and their instrumental role in regulating virulence and host specificity. In the past two decades, advances in genome sequencing have facilitated the sequencing of numerous genomes and the discovery of lineage-specific chromosomes in many economically significant plant pathogenic fungi (Kanapin et al., 2020; Henry et al., 2021; Wang et al., 2020; Zhang et al., 2020). The sequencing of accessory chromosomes from a multitude of formae species and their strains has illuminated potential origins of these chromosomes-whether they emerge from primary chromosomes through mechanisms such as misdivision or chromosomal rearrangements, or whether they are residues of ancient horizontal gene transfer events from other species. As the sequencing of fungal genomes continues to proliferate, comparative genomics between different formae speciales and their pathogenic races have provided substantial insights into the pathogenesis and host-specificity of *F. oxysporum*.

Despite *F. oxysporum* f. sp. carthami (Foc) being a catastrophic pathogen affecting safflower, a crucial oilseed crop, there has been a surprising absence of efforts to map the genomic information of its core and accessory genomes. In response to this knowledge gap, we conducted a systematic study, procuring 15 strains of Foc from the national depository. Our aim was to scrutinize their morphological traits and ascertain their pathogenic potential on host differentials. Our initial approach involved short-read sequencing of all the strains, subsequently establishing a phylogenetic relationship predicated on allelic variations. We then selected one virulent and one avirulent strain for Nanopore-based long-read sequencing. For the virulent strain, we further scaffolded the contig-level genome using Hi-C sequencing-based scaffolding protocols. Completion of chromosome level assembly of the genomes for both strains was achieved through the syntenic comparison of their scaffolds with the assembled and annotated genome of *F. oxysporum* f.sp. *lycopersici* (Ma et al., 2010). Upon completion, the estimated genome size for the virulent and avirulent strains stood at 46 Mb (comprising 12,902 coding genes) and 42 Mb (consisting of 10,991 coding genes), respectively. A comparative genomic analysis involving key gene families like effectors, secondary metabolites, carbohydrate-active enzymes, and mycotoxins revealed significant differences between the two genomes, potentially indicative of their respective pathogenic capabilities. We proceeded to infect a Fusarium wilt-resistant and -susceptible safflower line with the virulent strain, followed by an RNA-seq procedure to generate a mixed transcriptome. The fungus specific reads were segregated by mapping the RNA-seq reads to the assembled Fusarium genome and were further analyzed to identify the differentially expressed genes. In summary, our work presents the full-length genomes of a virulent and avirulent strain of Foc, their comparative differences, and the correlation between these differences and pathogenesis. This has profoundly enhanced our understanding of host-pathogen interactions and the adaptation strategies of these fungi. The insights gleaned from our study have the potential to guide the development of novel strategies to control these pathogens, thereby mitigating the effects of the diseases they instigate.

## Material and methods

### Fungal isolates

The *F. oxysporum* strain ‘FOC-IARI-5175’ used in this study was sourced from the Division of Plant Pathology, Indian Agricultural Research Institute (IARI), New Delhi. We procured an additional fifteen isolates of *F. oxysporum* f. sp. carthami (Foc) from ICAR-National Bureau of Agriculturally Important Microorganisms (NBAIM), India (Supplementary Table 1). All fungal strains were cultured, and single colonies were utilized to maintain their subcultures on potato dextrose agar (PDA) media, supplemented with cefotaxime (100 mg/L). These cultures were incubated for 3-4 days at a temperature of 28 ± 1°C in a New Brunswick Scientific incubator, USA.

### Morphological characterization

Morphological variations among Foc strains were evaluated based on several factors, including colony morphology, spore size, sporulation, and germination rate. These assessments were conducted on 7-day old Foc strains cultured on petri dishes with potato dextrose agar (PDA; Merck, Darmstadt, Germany). An approximately 5 mm piece of the actively growing colony from each isolate was placed on PDA plates and incubated at 27°C. After 7 days, each Foc colony was inspected for its colour, margin, and saltation, following the methodology described by Santos et al. (2019). Fungal spores were harvested from the PDA plates by adding 2 ml of sterile water, then filtering through sterile gauze to eliminate hyphal debris. The resulting conidia concentration was adjusted to 10^6^ spores/ml using a standard haemocytometer. All culture suspensions were examined on sterile slides in three replicates under a microscope (Olympus) at 40X magnification. Sporulation, length, and width of five conidia per slide from all strains were measured across three microscopic fields. For the germination test of conidia, a sucrose water suspension (10 mg/ml) was added to a 1.5 ml microcentrifuge tube (MCT) containing the spore suspension, and then incubated at 27°C for 24 hours. Following this period, visible germination structures were captured, and the germination rate was recorded.

### Measurement of growth rate

To measure the growth rate, a 5 mm piece of actively growing culture was excised from PDA plates and inoculated into 20 ml of Potato Dextrose Broth (PDB) medium supplemented with cefotaxime (50 mg/L). The inoculated medium was placed in an incubator shaker and maintained at a temperature of 26±2°C, with a speed of 200 rpm, for a duration of 3-4 days, followed by counting of the spores. Spores, with a density of 1 x10^4^ per ml, were then inoculated at the centre of fresh PDA plates, in triplicates. The diameter of the growing colony was measured, taking five readings each time, on the 2^nd^, 4^th^, 6^th^, and 8^th^ days post inoculation. The growth rate was studied and represented in mm/day for each Foc strain.

### Pathogenicity testing of strains

In this study, the pathogenicity of Foc isolates was tested using Foc-susceptible (PI-199897) and Foc-resistant (Sharda) safflower accessions. Safflower seeds from both accessions were surface sterilized and germinated under photoperiodic conditions of 16 h of light and 8 h of darkness, with a light intensity of 135 mmol/m^2^/s, at a temperature of 25±1°C. Each Foc strain was inoculated in Potato Dextrose Broth (PDB) medium and incubated in shaker incubators at 25±1°C for 2-3 days at 200 rpm. Resulting cultures were filtered through muslin cloth, and the spore count was adjusted to 0.75 x 10^6^ spores/ml using a haemocytometer. Fifteen-day-old healthy safflower seedlings were then immersed in the conidial suspension for 40 hours, after which the spore suspension was replaced with sterile water. A detectable phenotype emerged after 15 to 17 days. Depending on their disease reactions to PI-199897 (Foc-susceptible) and Sharda (Foc-resistant) accessions, strains were classified into different pathogenicity groups.

### Genomic DNA extraction and library preparation

For the extraction of genomic DNA, each isolate was cultured in 50 ml of Potato Dextrose Broth (PDB) in a 250 ml conical flask, followed by incubation at 26±1°C for 3-4 days. Fungal mycelia were collected by filtering the culture through eight layers of muslin cloth. High-quality genomic DNA was then extracted using the Quick-DNA Fungal/Bacterial Miniprep Kit from Zymo Research, following the manufacturer’s instructions. The integrity and concentration of the genomic DNA were estimated on a 0.8% agarose gel. The genomes of a virulent strain (Foc-IARI-5175) and an avirulent strain (F-00845) were sequenced using Nanopore sequencing technology to develop reference genomes. Additionally, all other Foc strains in this study were sequenced through Illumina’s short-read sequencing technology. For Nanopore sequencing, libraries were prepared from high quality DNA using the ‘Ligation sequencing kit 1D’ following the manufacturer’s instructions (Oxford Nanopore, UK). In brief, around 2 μg of high molecular weight DNA was repaired using the ‘NEBNext FFPE DNA Repair mix’ and the ‘Ultra ll End-prep Enzyme mix’; subsequently, the adapter mix was ligated to the repaired DNA using the ‘NEBNext Quick T4 DNA Ligase’. At the end of each step, DNA was cleaned with the ‘AMPure XP beads’ (Thermo Fisher Scientific, USA). The quality and quantity of the DNA libraries were determined with a Nanodrop spectrophotometer. DNA libraries were sequenced on the MinION device using the MinION Flow Cells R 9.4.1 (Oxford Nanopore, UK).

For Illumina sequencing, paired-end libraries of 250 bp insert size were developed using the NEBNext Ultra II FS DNA library prep kit for Illumina, as per the instructions provided by the manufacturer. The yield of the constructed library was quantified using a Qubit fluorometer (Invitrogen), and the size was determined using a Bioanalyzer (Agilent Technologies Inc., USA) with a DNA HS chip. The developed illumina librarier were sequenced on a HiSeq-2000 platform.

For the IARI-5175 strain, additional Hi-C sequencing was performed to achieve chromosomal-level ordering of scaffolds. The Hi-C library for fungal samples was prepared using the Arima-HiC kit (Arima Genomics, Inc.; Cat # A510008) following the manufacturer’s protocols. In brief, the fungal samples were homogenized into a fine white powder under liquid Nitrogen and subsequently crosslinked. The crosslinked samples were then subjected to the Arima-HiC protocol, which uses a combination of enzymes to digest chromatin inside the nuclei. The processed DNA was then sheared to generate fragments ranging from 200 to 650 bp, which was used to generate Illumina sequencing libraries. Finally, the Illumina NovaSeq sequencer was used to sequence the libraries using the 150 bp PE mode to generate 3 Gb of high-quality data.

### Genome Assembly

Base-calling and quality filtering of the sequencing reads generated from the Minion sequencer were performed using the Albacore software (v2.5.11; https://github. com/Albacore). Quality control (QC) passed reads with length greater than 500 bp were used for contig generation. The assembly of reads into contigs was carried out using the CANU (V1.4) software (Koren et al., 2017), which utilizes a hierarchical assembly pipeline. The process of generating contigs was divided into three stages: correction, trimming, and finding the unitiggs. Quality control of the generated contigs was carried out, and any reads less than 500 bp were filtered out. The contigs were then arranged into scaffolds using SALSA (Ghurye et al, 2019) for Hi-C-based scaffolding. The Hi-C-generated reads were mapped onto the contigs, and scaffolds were generated using SALSA. This process allowed for the spatial arrangement of the contigs according to their positions in the genome, providing a more accurate representation of the genome’s structure.

### Phylogenomic analysis

Phylogenomic analysis was conducted on thirteen Foc strains in order to understand and reconstruct the evolutionary relationship among these strains. Two closely related formae speciales, *F. oxysporum* f. sp. *ciceris* (SRA ID-SRR837400) and *F. oxysporum* f. sp. *lycopersici* (SRA ID-SRR7690004), as well as a non-pathogenic strain, *F. oxysporum* FO47 (SRA ID-SRR11523115), were included as outgroups. Whole genome sequencing reads were mapped onto the reference genome of *F. oxysporum* f. sp. lycopersici using the Bowtie assembler. Single Nucleotide Polymorphisms (SNPs) were then filtered from all strains, with a minor allele frequency (MAF) cut-off of 0.05. Following this, a neighbor joining (NJ) tree was constructed using the maximum likelihood method with the MEGA software. The tree obtained in Newick format was then converted to a graph using the Interactive Tree of Life (iTOL) software for visualization and further analysis.

### Analysis of repeat element and gene prediction

In order to identify the repeat sequences in the Foc genome, a library encompassing all known repeat sequences was compiled. This was accomplished through the creation of a *de novo* library from the assembled genome utilizing the RepeatModeler pipeline (http://www.repeatmasker.org/ RepeatModeler/). The final repeat database was produced using several software programs, including RECON (Bao and Eddy, 2002), RepeatScout (Price et al., 2005), Tandem Repeat Finder (Benson and Dong, 1999), and NSEG (ftp://ftp.ncbi.nih.gov/pub/seg/nseg/). This newly generated database/library was then utilized to mask repeat elements in the assembled genome using the RepeatMasker software. This process served to mark the locations of repeat elements within the genome sequence, making these areas easily identifiable for downstream analyses. Once the repeat sequences were masked, protein coding genes were identified in the repeat-masked genome using the Augustus software (Stanke & Morgenstern, 2005).

### Comparative genomics with Fol and Identification of accessory genome

The genes predicted in the Foc genome were compared with those of *F. oxysporum* f. sp. lycopersici (Fol) in order to identify the core genome, which is the genomic region displaying synteny with Fol, and the accessory genome, which consists of unique sequences present only in Foc strains. To conduct this comparison, the total protein sequences of both the assembled genomes were subjected to a BlastP analysis with an e-value threshold of 1e-05. After the BlastP analysis, the MCScan software was used to identify syntenic regions between the two genomes. The threshold used for the identification of a region as a syntenic stretch was set at a minimum of five genes in a row.

### Prediction of candidate secretory effector proteins, carbohydrate active enzymes (CAZymes) and secondary metabolite gene clusters

The search for putative secretory effector proteins was conducted by screening the predicted protein coding genes in the IARI-5175 and F-00845 strains using the EffectorP 3.0 software. Predicted effector proteins were further scrutinized through sequential screening using various online tools like SignalP 5.0, TargetP, TMHMM 2.0, and PredGPI. These tools provided predictions about presence and location signal peptide, subcellular localization, transmembrane helices, and GPI (glycosylphosphatidylinositol) anchor signals. The effector proteins with signal sequence and without predicted transmembrane domain, subcellular localization signal, and GPI anchor signals were retained. The identification and annotation of CAZymes (Carbohydrate-Active enZymes), enzymes that are involved in the metabolism of carbohydrates, was performed using the dbCAN online tool. Finally, to identify secondary metabolite gene clusters, the antibiotics and secondary metabolite analysis shell (antiSMASH v6.0) online tool was utilized.

### RNA isolation and preparation of transcriptome libraries

Safflower accessions susceptible (PI-199897) and resistant (Sharda) to the Foc-IARI-5175 strain were used to prepare tissue for transcriptome analysis. The safflower seedlings were stressed, and the infected roots were harvested 7 days post-infection (dpi). Foc mycelia were also gathered to serve as a control in this study. Two biological replicates were gathered for each sample. RNA isolation was performed using TRIzol reagent from both infected and control samples. Around 500 mg of tissue was ground to a fine powder in liquid nitrogen. This powder was then thoroughly mixed with 5 ml of TRIzol reagent, allowed to thaw at room temperature, and then transferred to a sterile centrifuge tube. The mixture was vortexed and allowed to rest for 5 more minutes at room temperature. Next, 1/3rd volume of chloroform was added to the mixture and vortexed thoroughly, followed by centrifugation at 4°C, 12000 x g, for 10 min. The aqueous layer was carefully transferred to a fresh falcon tube and equal amounts of Phenol (pH 4.3): Chloroform: Isoamyl alcohol (25:24:1) mixture were added. This homogenate was mixed again by vortexing followed by centrifugation. After centrifugation, the aqueous layer was transferred to a sterile falcon tube and purified via two sequential chloroform extractions. The purified aqueous layer was mixed with an equal volume of pre-chilled isopropanol and incubated in −20°C for 2 hours for precipitation. The precipitated RNA was collected via centrifugation at 4°C, 12000 x g, for 15 min. The resultant RNA pellet was washed with 70% ethanol and centrifuged at 25°C, 12000 x g for 5 min to remove salts. The RNA pellet was allowed to air dry and then re-dissolved in 100 µl of nuclease-free water. The RNA was then quantified using a spectrophotometer and its integrity was checked on a 1.2% denaturing agarose gel. The samples were re-precipitated in ethanol and stored at - 80°C until further processing. For generation of transcriptome libraries, a total of 10 ng of purified RNA was used to generate paired end libraries using the NEBNext® Ultra II Directional RNA Library Prep Kit, according to the manufacturer’s directions. The libraries were initially quantified using a Qubit fluorometer and diluted to 1 ng/µl. The size of each library was determined using a Bioanalyzer with a DNA HS chip.

### Transcriptome sequencing and data analysis

All library preparations were sequenced using the Illumina NovaSeq6000 platform. The raw read quality was evaluated using FastQC, followed by the removal of adapter sequences and low-quality reads utilizing Trimmomatic software with parameters set as TruSeq3-PE-2.fa:2:30:10 LEADING:3 SLIDINGWINDOW:4:15 TRAILING:3 MINLEN:60. The remaining high-quality reads were then aligned to the reference genome of the IARI-5175 strain, developed in this study, using the STAR aligner (Dobin et al., 2013). According to the position information for each gene, the number of reads covering each gene was counted using the STAR aligner. Reads aligning to multiple regions in the genome or reads with low alignment quality were filtered out. Gene expression levels were estimated using the measure “fragments per kilobase per million mapped fragments” (FPKM). This measure takes into account the effects of both sequencing depth and gene length for the read counts. Differentially expressed genes were identified by DESeq2 v1.30.1 (Love et al., 2014) using Bioconductor (Gentleman et al., 2004) in R (R Core Team, 2019).

## Results

### Pathogen identification, morphological characterization and pathogenicity screening

The acquired strains were initially subcultured on Potato Dextrose Agar (PDA) plates for single colony isolation and subsequently verified using a Foc-specific Sequence Characterized Amplified Region (SCAR) marker (Singh & Kapoor, 2018). The presence of a 213 base pair band affirmed the homogeneity and purity of the procured Foc strains (Supplementary Figure 1A). Examination of the colony margin curvature revealed either a circular or wavy morphology (Table 1). Upon closer scrutiny, we observed that colonies for four of the strains (F-00945, F-00946, F-02217, and F-02220) exhibited a dull white hue, whereas the colonies of the remaining strains (F-02003, F-00841, F-00845, F-00924, F-00933, F-00934, F-00938, F-00940, F-02218, F-02219, and F-02221) bore an orangish coloration. The growth rates varied across the strains, spanning from 4.46 to 6.98 mm/day (Figure 1A), with F-00933 displaying the most rapid growth and F-02019 growing at the slowest rate. Further microscopic analysis revealed that the length of the microconidia oscillated between 3.75 µm (in F-02218) and 6.03 µm (in F-00946), while the width fluctuated from 1.03 µm (in F-02217) to 1.41 µm (in F-00938). Of all the strains, IARI-5175 exhibited high sporulation and recorded the highest germination percentage (78.75%). Conversely, strain F-00946, which demonstrated low sporulation, also had the lowest spore germination percentage at 28% (Table 1).

**Figure 1:**
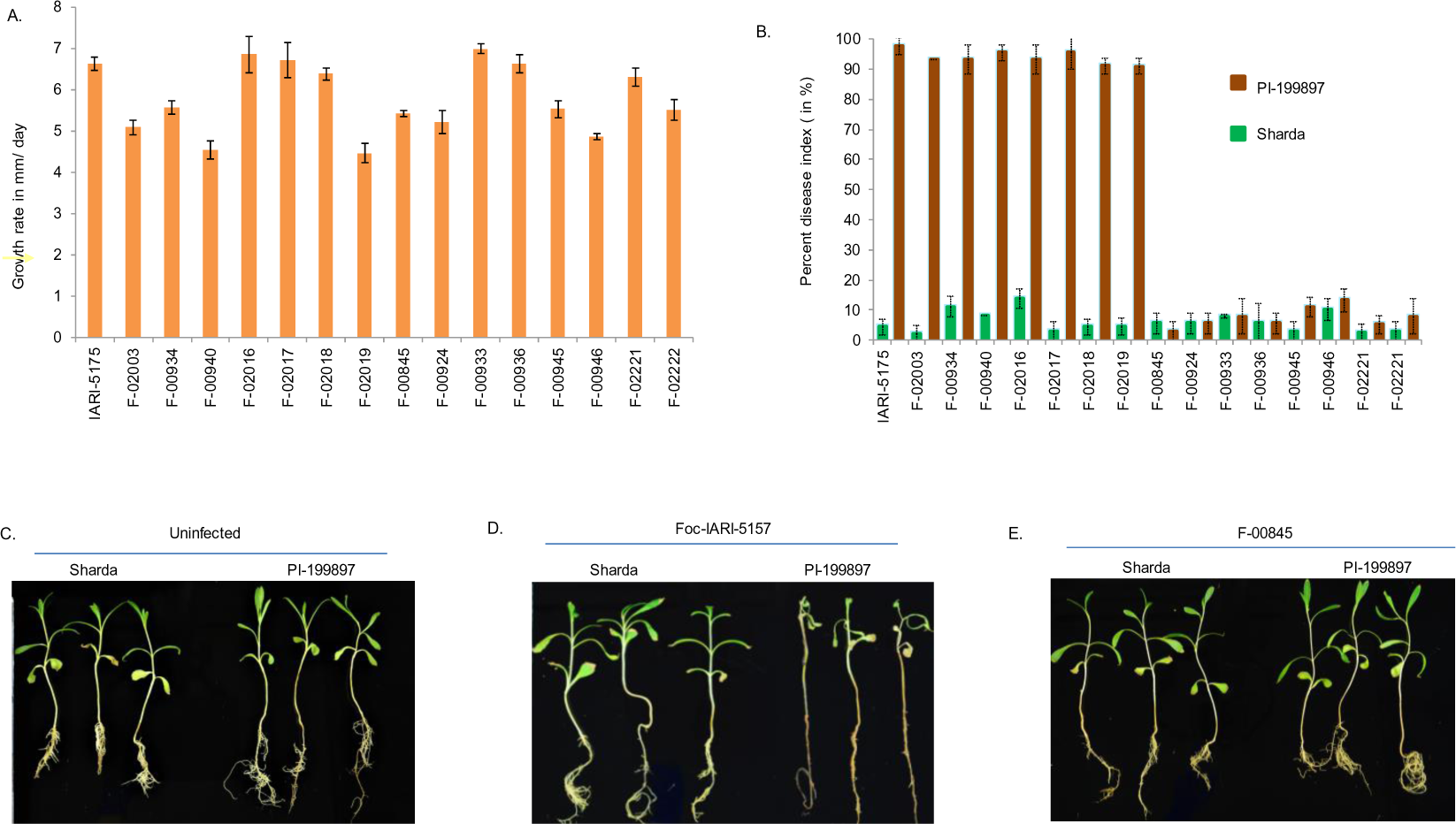
Confirmation and pathogenicity testing of Foc strains. A) Growth rate measurement of Foc strains in mm/day. B) Percentage disease incidence (PDI) for phenotyping of strains in Foc susceptible and Foc resistant safflower accession. C) Uninfected control Safflower accessions. D) Foc-IARI-5175 showing avirulent phenotype with both Sharda and PI-199897. E) Foc-00845 strain showing avirulent and virulent phenotype with Sharda and PI-199897 respectively.

**Table 1:** Growth rate, colony morphology, PDI and microconidial dimensions of Foc strains cultured on potato dextrose agar medium.

| S. No. | Foc strain | Growth rate (mm/day) | Colony margin | PDI ( in %) |  | Microconidial dimensions (µm) |  |
| --- | --- | --- | --- | --- | --- | --- | --- |
|  |  |  |  | Sharda | PI-199897 | Length | cc |
| 1 | IARI-5175 | 6.628 | Circular | 4.4 | 97.7 | 4.07 | 1.24 |
| 2 | F-02003 | 5.08 | Wavy | 2.2 | 93.3 | 3.87 | 1.30 |
| 3 | F-00934 | 5.57 | Circular | 11.06 | 93.3 | 4.22 | 1.18 |
| 4 | F-00940 | 4.53 | Wavy | 8.3 | 95.5 | 4.39 | 1.3 |
| 5 | F-02216 | 6.85 | Circular | 13.73 | 93.3 | 5.08 | 1.40 |
| 6 | F-02217 | 6.72 | Wavy | 2.76 | 95.5 | 4.04 | 1.02 |
| 7 | F-02218 | 6.38 | Circular | 4.4 | 91.0 | 3.75 | 1.27 |
| 8 | F-02219 | 4.46 | Circular | 4.56 | 90.9 | 4.55 | 1.21 |
| 9 | F-00845 | 5.422 | Wavy | 5.53 | 2.76 | 4.49 | 1.28 |
| 10 | F-00924 | 5.2 | Circular | 5.53 | 5.53 | 4.63 | 1.18 |
| 11 | F-00933 | 6.98 | Circular | 7.9 | 7.9 ± 5.9 | 4.02 | 1.28 |
| 12 | F-00936 | 6.62 | Circular | 5.53 | 5.53 | 3.88 | 1.33 |
| 13 | F-00945 | 5.52 | Wavy | 2.27 | 10.9 | 4.21 | 1.29 |
| 14 | F-00946 | 4.86 | Wavy | 10.16 | 13.23 | 6.03 | 1.29 |
| 15 | F-02221 | 6.3 | Wavy | 2.36 | 5.13 | 4.40 | 1.27 |
| 16 | F-02222 | 5.504 | Circular | 2.76 | 7.9 | 4.3 | 1.47 |

In an attempt to identify any correlation between growth differences and the strains’ pathogenicity, we carried out a pathogenicity test utilizing a hydroponics-based method (Kukreja et al., 2018). To span the spectrum of avirulence, virulence, and hypervirulence, we evaluated two safflower accessions: a susceptible one (PI-199897) and a more tolerant one (Sharda) (Singh et al., 2022). Strains that infected both the resistant and susceptible lines were classified as hypervirulent, while those that failed to infect either line were designated as avirulent. Strains that could infect the susceptible safflower line but not the tolerant one were deemed virulent. On the basis of the ratio of dead to surviving plants following 16 days of infection, the Foc strains F-02003, F-00934, F-00940, F-02216, F-02217, F-02218, F-02219, and IARI-5175 demonstrated virulence or compatible reactions with the PI-199897 accession (Supplementary Figure 1I-O). The resistant accession, Sharda, exhibited minor symptoms with these strains, yet none of the plants succumbed, thereby indicating avirulence. Conversely, strains F-00845, F-00924, F-00933, F-00936, F-00945, F-00946, F-02221, and F-02222 were incapable of infecting either the susceptible or resistant accessions and were thus labelled as exhibiting avirulence or incompatible reactions (Supplementary Figure 1B-H). None of the strains demonstrated compatible interactions with both the susceptible and resistant accessions, which implies that none were hypervirulent. Taken together, out of the 16 strains tested, F-00845, F-00924, F-00933, F-00936, F-00945, F-00946, F-02221 and F-02222 were determined to be avirulent, whereas F-02003, F-00934, F-00940, F-02216, F-02217, F-02218, F-02219, and IARI-5175 were categorized as virulent (Supplementary Figure1B-O).

### Phylogenomic analysis

To decode the molecular phylogeny and ascertain the evolutionary genesis of the *F. oxysporum* f. sp. *carthami* (Foc) strains examined in our study, we retrieved genome-wide Single Nucleotide Polymorphisms (SNPs) from the contig-level assembled genomes of these strains. However, we could only successfully extract high-quality SNPs from a subset of 13 strains, consisting of eight virulent and five avirulent. Consequently, these thirteen Foc strains served as the foundation for constructing our phylogenetic tree. Additionally, we included three distinct forms of *F. oxysporum* (f. sp. *ciceris*, f. sp. *lycopersici*, and FO47) as outgroups in our analysis, and derived a neighbour-joining tree using the maximum likelihood method (Figure 2). In the resulting phylogenetic tree, all thirteen Foc strains coalesced into a single clade situated adjacent to the outgroup clade. This configuration suggests a probable monophyletic origin for all Foc strains used in our study. Interestingly, *F. oxysporum* f. sp. *ciceris* emerged as the closest relative within the outgroup, thus indicating its intimate genetic affinity with Foc. Within the Foc clade, avirulent strains (F-00936, F-00945, F-02221, F-00845, and F-00946) were found scattered among four virulent strains (F-02217, F-02218, F-02219, and F-00940). In contrast, four other virulent strains (F-00934, F-02216, IARI-5175, and F-02003) formed a distinct clade, which implies that these specific Foc strains may have procured their virulence via individual evolutionary paths. This observation underscores the potential for significant genetic variation within a single pathogen species, which can engender disparities in pathogenicity and host specificity.

**Figure 2:**
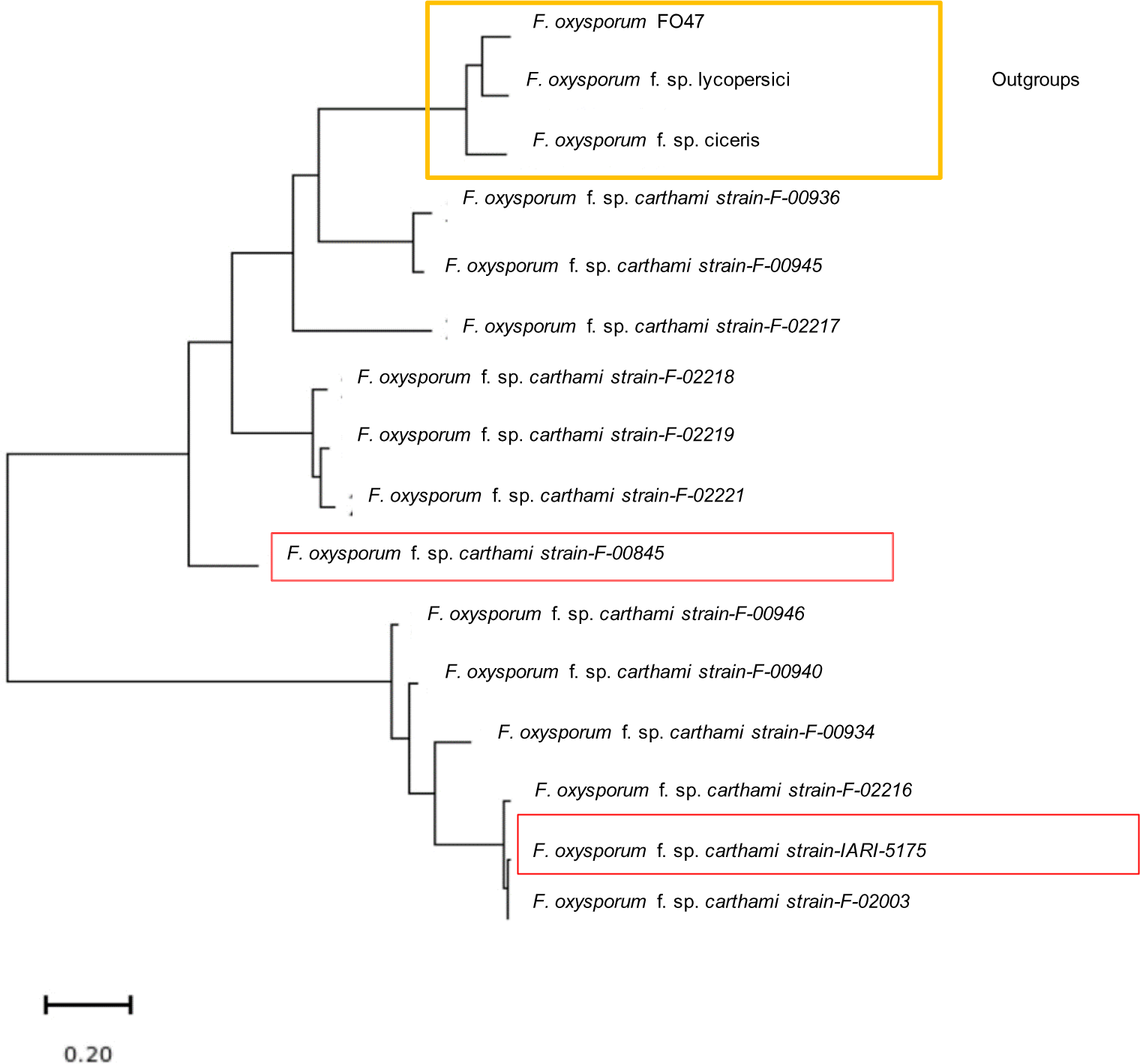
Phylogenetic placement of Foc strains with closely related formae speciales of *F. oxysporum*. Phylogeny of Foc strains was inferred from neighbor joining tree obtained by maximum likelihood method. SNPs were extracted from all genomes under study and the tree was constructed through MEGA software. Strains highlighted in orange box were included as outgroups and strains highlighted in red boxes are selected virulent (IARI-5175) and avirulent strain (F-00845) for comparative genomic analysis.

### Genome assemblies of representative virulent and avirulent Foc strains

For a comprehensive comparative genomic study, we chose Foc strains IARI-5175 and F-00845 as prototypical examples from virulent and avirulent pathogenic groups, respectively (Figure 1D and 1E). Utilizing Illumina short read sequences, we conducted a K-mer analysis, which estimated the sizes of the assembled genomes to be 57.7 Mb for the virulent Foc strain (IARI-5175) and approximately 52.3 Mb for the avirulent Foc strain (F-00845). To further delve into the genomic difference between the virulent and avirulent strains, we utilized long-read Nanopore-based sequencing for IARI-5175 and F-00845. This method produced a total of 46,473,015 and 42,479,514 high-quality raw reads, which were assembled into 157 and 72 contigs for IARI-5175 and F-00845, respectively. Further refinement via hierarchical scaffolding, using the HiC sequences, resulted in 88 and 23 scaffolds for IARI-5175 and F-00845, respectively, culminating in a combined length of approximately 46.73 Mb and 42.48 Mb. The assembled genomes of IARI-5175 and F-00845 house 12,902 and 10,991 protein-coding genes respectively, with total lengths of 4,882,719 bp and 8,380,456 bp. Table 2 provides an in-depth presentation of the statistics related to the genome assembly of IARI-5175 and F-00845.

**Table 2:** Genome assembly statistics of virulent (IARI-5175) and avirulent (F-00845) Foc strains at contig and scaffold level.

| S. No. | Parameter | IARI-5175 | F-00845 |
| --- | --- | --- | --- |
| **Contig level assembly** |  |  |  |
| 1 | Sequence count | 172 | 57 |
| 2 | Size of assembly | 46473015 | 42479514 |
| 3 | Longest contig | 2365047 | 4374376 |
| 4 | N50 | 785227 | 1773145 |
| 5 | L50 | 16 | 7 |
| 6 | GC percentage | 47.98 | 48.39 |
| **Scaffold level assembly** |  |  |  |
| 7 | Sequence count | 88 | 23 |
| 8 | Size of assembly | 46735215 | 42481214 |
| 9 | Longest contig | 6762594 | 6040854 |
| 10 | N50 | 4521379 | 4391618 |
| 11 | L50 | 4 | 4 |
| 12 | GC percentage | 47.98 | 48.39 |
| 13 | Protein coding genes | 12902 | 10991 |
| 14 | Total length of Protein coding genes | 4882719 | 8380456 |

### Comparative genomics analysis with Fol resulted in chromosomal level assembly and identification of accessory genome in Foc chromosomal level assembly, comparative genome analysis and identification of accessory genome in Foc

The genomic sequences of IARI-5175 and F-00845 were aligned with the reference genome of Fol4287, a well-characterized member of the *F. oxysporum* species complex (FOSC), (accession number AAXH00000000.1). This alignment process involved performing pairwise BLASTp comparisons between Fol and the individual Foc strains, resulting in a dot plot (Figure 3A and 3C). Following this alignment with the reference genome, the scaffolds for IARI-5175 and F-00845 were assembled into 11 core chromosomes (Figure 4A and 4B). As expected, no syntenic regions within the assembled genomes of both Foc strains corresponded to the four accessory chromosomes (3,6,14, and 15) of Fol4287. The size of the core genome in IARI-5175 and F-00845 was observed to be 40.4 and 39.8 Mb, respectively. Certain scaffolds of IARI-5175 (∼6.2 Mb) and F-00845 (∼2.6 Mb) did not align with the Fol4287 genome, thereby leading to their designation as lineage-specific genomic regions. Specifically, two lineage-specific scaffolds in strain IARI-5175 exceeded 0.75 Mb in size and were therefore considered as lineage-specific chromosomes. These included scaffold u000000273 of 1.34 Mb and u000000192 of 1.33 Mb (Figure 4A). Similarly, in avirulent Foc strain F-00845 two scaffolds tig0000021 and tig0000041 of size 1.0 Mb and 0.867 Mb were designated as lineage specific chromosomes (Figure 4B). In an attempt to identify the genes uniquely present in IARI-5175 and F-00845, we compared the 17,447 unique genes of Fol4287 with 12,902 genes of IARI-5175 and 10,991 genes of F-00845 (Figure 3B and 3D). This comparison revealed that approximately 94.05% (12,135 genes) of IARI-5175 and 94.04% (10,337 genes) of F-00845 were similar to Fol4287. The number of unique genes in IARI-5175 and F-00845 with respect to the Fol4287 genome was 767 and 654, respectively. Notably, all these unique genes were located within the lineage-specific region of their respective genomes. Comparison of genomes between the two Foc strains revealed that 407 genes were uniquely present in the accessory genome of Foc-IARI 5175.

**Figure 3:**
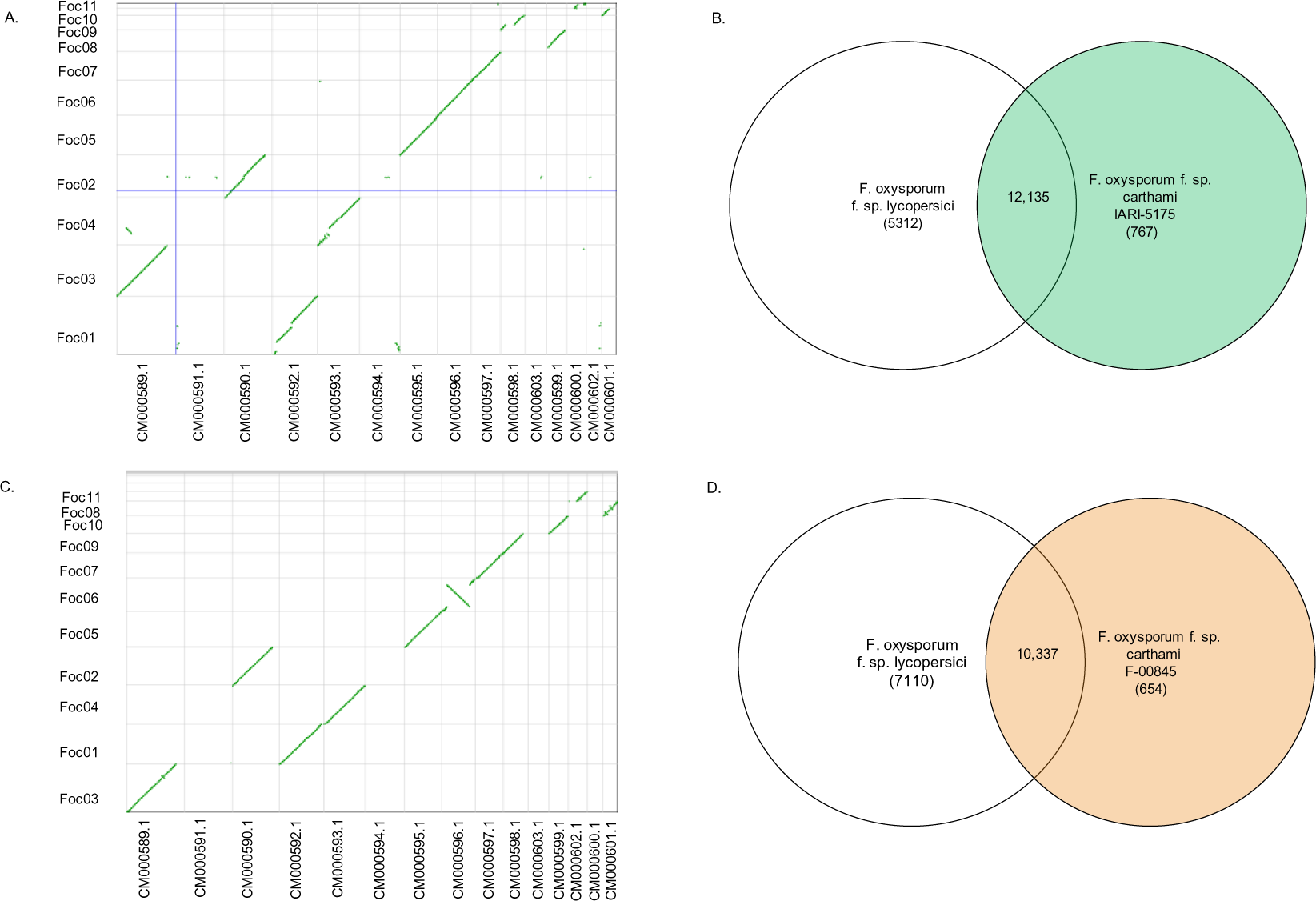
Comparative genomic analysis of Foc strains IARI-5175 and F-00845 with *F. oxysporum* f. sp*. lycopersici*. A) Dotplot of pairwise BLASTp alignment between IARI-5175 and Fol showing chromosome correspondence between two genomes. B) Venn diagram showing unique and shared number of genes between IARI-5175 and FOL 4287. C) Dotplot of pairwise BLASTp alignment between F-00845 and Fol showing chromosome correspondence between two gen omes. D) Venn diagram showing unique and shared number of genes between F-00845 and FOL 4287.

**Figure 4:**
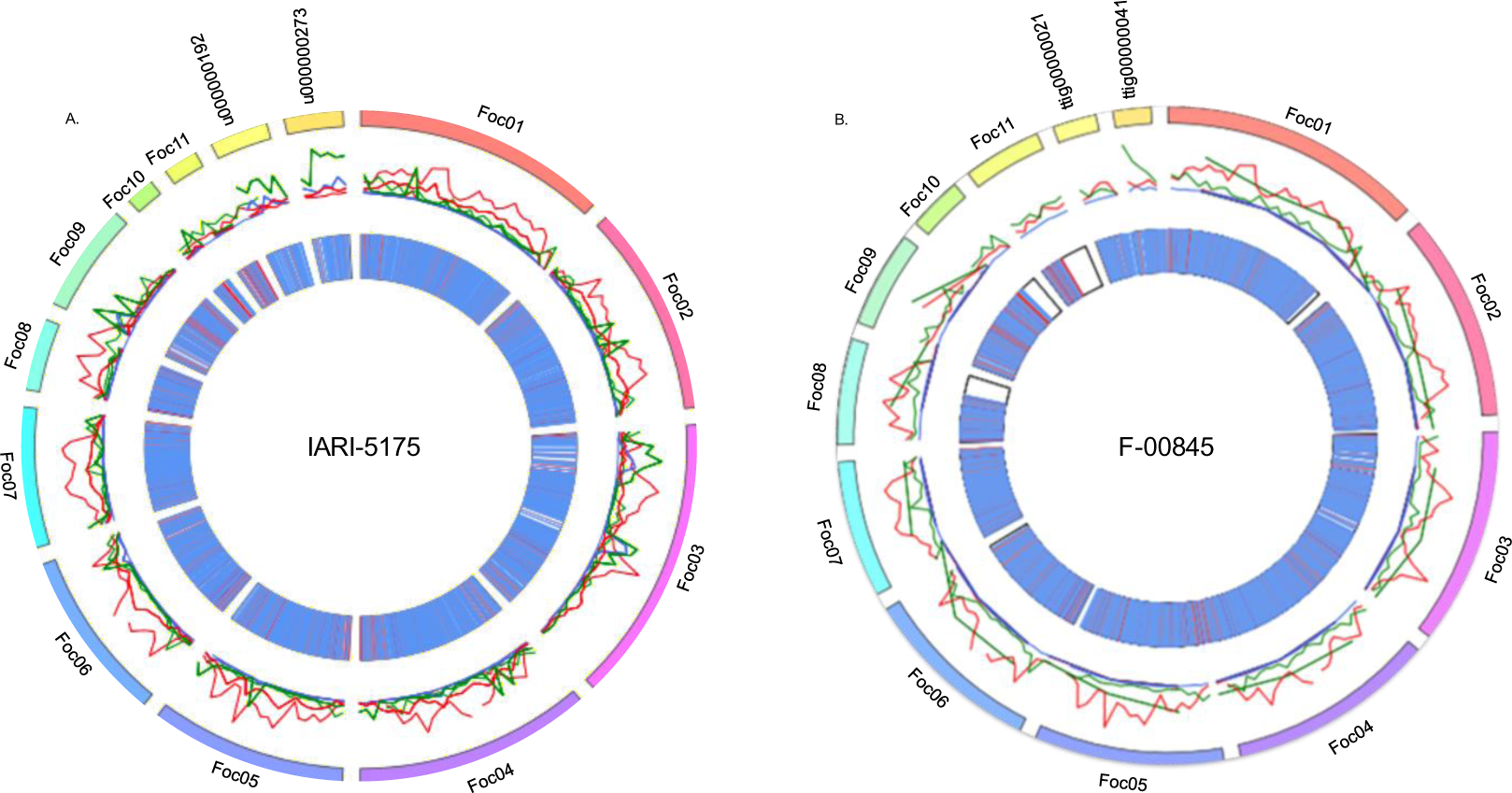
Visualization of genomes of representative virulent and avirulent Foc strains. A) core and lineage specific regions of IARI-5175 (virulent) strain wherein different tiers from outside to inside represents chromosomal designation in accordance with Fol and two lineage specific chromosomes, repeat elements and genes encoding for putative effector proteins. B) A) core and lineage specific regions of F-00845 (avirulent) strain wherein different tiers from outside to inside represents chromosomal designation in accordance with Fol and two lineage specific chromosomes, repeat elements and genes encoding for putative effector proteins.

### Repeat elements

For the prediction of repeat elements in both assembled genomes, we generated a *de-novo* repeat library consisting of 412 consensus repeat sequences. This newly generated library was then utilized to predict repeat elements. The assembled genome of IARI-5175 exhibited a higher proportion of repeat sequences (∼18%) compared to that of F-00845 (∼8%). Given that these repeat sequences are predominantly AT-rich, the genome of IARI-5175 consequently manifested a lower GC content (47.98%) compared to F-00845 (48.40%). In terms of classified retroelements, LTR elements constituted the most abundant portion in the genomes of both strains [IARI-5175 (1.28%) and F-00845 (0.28%)]. For IARI-5175, the remaining retroelements were found in the following decreasing order of abundance: LINEs, Ty1/Copia, and Gypsy/DIRS1. Conversely, the genome of F-00845 lacked Ty1/Copia but exhibited a greater abundance of Gypsy/DIRS1 compared to LINE elements. Among the DNA transposons, hobo-Activator and Pogo type were the major contributors. IARI-5175 additionally contained PiggyBac type DNA transposons, which were absent in F-00845. The genomes of IARI-5175 and F-00845 contained 11.23% and 2.21% interspersed repeats, and 0.7% and 0.46% tandem repeats, respectively (Supplementary Table 2). In addition to the classified repeats, the genomes of IARI-5175 and F-00845 also contained 4.91% and 1.83% unclassified repeats respectively (Supplementary Table 2 and Figure 4B).

### A comparison of the potential effectors between the virulent and avirulent strain

In an effort to uncover the potential basis of virulence mediated by effector proteins, we deployed an elimination pipeline to predict effectors from both virulent and avirulent Foc strains (Figure 5A). Out of the 12,902 proteins from the IARI-5175 strain, 5,156 proteins were tentatively identified as effectors via the application of EffectorP (Sperschneider & Dodds, 2022) (Figure 4A). Likewise, out of 10,991 proteins from the F-00845 strain, 1,940 effectors were pinpointed (Figure 4B). Subsequently, we sought out the secretory effectors among the predicted ones by processing them with the help of SignalP. By employing TargetP, TMHMM, and PredGPI software, we managed to identify and subsequently exclude those effectors that targeted organelles, plasma membranes, and cell walls based on their specific signature sequences. In the end, we identified 226 and 101 effectors from the IARI-5175 and F-00845 strains, respectively (Figure 5A). The core genomes of IARI-5175 and F-00845 harboured 215 and 98 effectors, respectively, while the lineage-specific genomes contained 11 and 3 effectors, respectively. As expected, the density of effectors in the IARI-5175 genome (4.83 effectors/Mb) was 2.03 times higher than in the F-00845 genome (2.37 effectors/Mb). Among all chromosomes, Chromosome-5 of IARI-5175 and Chromosome-9 of F-00845 boasted the highest numbers of effectors, numbering 32 and 17, respectively (Table 3). When we compared the effectors of IARI-5175 and F-00845, we found only 20 to be common, while 206 were unique to the virulent strain (Figure 5B). All of these common effectors were located within the core genome of IARI-5175 and F-00845. Of note, nearly 95% (195) of the 206 effectors unique to IARI-5175 were present in the core genome, while the remaining ∼5% (11) were situated in the lineage-specific genomic portion (Figure 5A)

**Figure 5:**
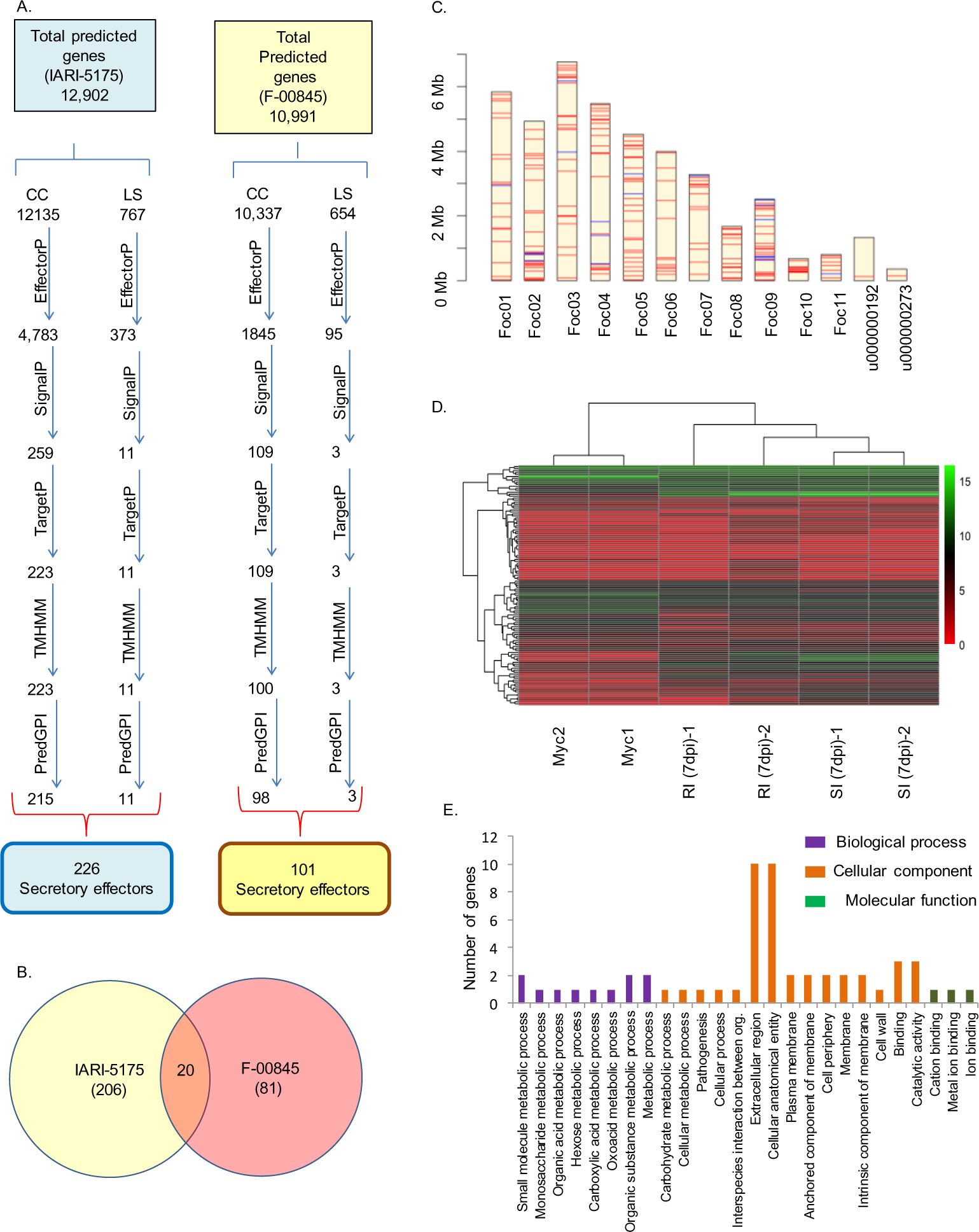
Prediction of secretory effectors in Foc and their characterization. A) Diagrammatic representation of pipeline used to identify candidate secretory effector proteins. B) Venn diagram of predicted genes coding for effector proteins in IARI-5175 and F-00845 strains. C) Chromosome wise gene block arrangement of effectors in IARI-5175. D) Clustering and heatmap of effectors present uniquely in IARI-5175 showing expression of effectors in control mycelia, 7day post infection (dpi) susceptible infected and 7 dpi resistant infected safflower accessions (2 replicates for each treatment). E) GO enrichment analysis of unique effectors in IARI-5175.

**Table 3:** Details of chromosome-wise distribution of effectors and effector density in Foc strains IARI-5175 and F-00845.

| Chromosome ID | Number of effectors in IARI-5175 | Effector density | Number of effectors in F-00845 | Effectors density |
| --- | --- | --- | --- | --- |
| Foc01 | 18 | (3.08/ Mb) | 10 | (1.65/ Mb) |
| Foc02 | 27 | (5.47/ Mb) | 10 | (2.12/Mb) |
| Foc03 | 26 | (3.84/ Mb) | 14 | (2.81/Mb) |
| Foc04 | 23 | (4.20/ Mb) | 8 | (1.67/Mb) |
| Foc05 | 32 | (7.02/ Mb) | 12 | (2.79/Mb) |
| Foc06 | 16 | (4/ Mb) | 7 | (1.70/Mb) |
| Foc07 | 13 | (4.06/Mb) | 3 | (0.96/Mb) |
| Foc08 | 14 | (8.75/Mb) | 3 | (1.25/Mb) |
| Foc09 | 29 | (11.6/Mb) | 17 | (8.09/Mb) |
| Foc-10 | 9 | (1.36/Mb) | 5 | (4.16/Mb) |
| Foc-11 | 8 | (10.12/Mb) | 9 | (5.29/Mb) |
| LS | 11 | (1.77/Mb) | 3 | (1.15/Mb) |

To elucidate the biological function of the unique effectors in the IARI-5175 strain, we conducted a Gene Ontology (GO) analysis. A striking 158 effector proteins fell under the GO category of “catalytic activity,” while 85 were classified as pertaining to “binding.” Interestingly, we noticed significant enrichment of GO terms such as “carbohydrate binding”, “chitin binding”, and “carbohydrate derivative binding,” which collectively encompassed 23 effectors. Although “lyase activity” and “isomerase activity” were also significantly represented, they included only four effectors each.

In terms of biological processes, a majority of unique effectors were associated with “metabolic processes” and its sibling GO terms. Notably, 57 and 56 proteins were related to the GO terms “Interspecies interaction between organisms” and “pathogenesis,” respectively. From the standpoint of cellular processes, all the enriched GO terms underscored the secretory nature of the unique effectors. For instance, 150 proteins corresponded to “extracellular regions,” defined as the space external to the cell’s outermost layer, while 192 proteins related to “cellular anatomical entity,” which denotes a substance produced by a cellular organism that surpasses the granularity of a protein complex (Figure 5E). A comprehensive list of GO terms for all analysed transcriptome groups is provided in Supplementary Table 5.

Simultaneously, we examined the expression of these effectors in Foc-infected resistant and sensitive safflower roots by conducting a whole transcriptome sequencing, using fungal mycelia as a control. During the compatible interaction, we observed differential regulation in 77 effectors as compared to the control. Among these, 49 effectors exhibited an increase in expression, while 28 were down regulated. In the incompatible interaction, 40 effectors showed differential regulation - 26 showed upregulation while 14 were down regulated.

Our ultimate goal was to discern the differences and similarities in the expression of effectors in resistant and susceptible hosts. Specifically, we categorized effectors into three classes: (i) Class-I, showing increased expression in the compatible interaction but reduced expression in the incompatible interaction; (ii) Class-II, exhibiting lower expression in the compatible interaction but higher in the incompatible interaction; and (iii) Class-III, with increased expression in both types of interactions. Our findings revealed 14 effectors in Class-I, 7 in Class-II, and 25 in Class-III. Effectors down regulated in both types of interactions were ruled out from further analysis as these are unlikely to be related to the host defence mechanism or pathogen virulence.

The findings presented in this work significantly contribute to our understanding of the biology and pathogenesis mechanisms of the Foc. The comprehensive Gene Ontology (GO) analysis provided insights into the broad range of biological functions of the identified unique effectors in the strain. A substantial number of these effectors were found to possess catalytic activity or were involved in binding, suggesting that they may be critical players in enzymatic processes or interactions with other molecules or cellular components, respectively. Interestingly, a significant number of these effectors were associated with carbohydrate binding and chitin binding, implying their potential role in interacting with host cell components and possibly facilitating pathogen infection. Furthermore, the unique effector proteins were predicted to have a significant role in metabolic processes, interspecies interactions, and pathogenesis. This implies that these effectors may be critical in the strain’s ability to interact with and infect host organisms, and perform metabolic activities that sustain its survival and growth. The work also pointed out the secretory nature of these unique effectors, which aligns with the general understanding that pathogens often secrete effector proteins to modulate host cell processes, facilitate infection, and evade the host’s immune response.

Through transcriptome sequencing, our study provided an insightful snapshot of how the expression of effectors change in response to infection in both resistant and sensitive hosts. Such understanding is pivotal in revealing the molecular dynamics of host-pathogen interactions. The identification of the three classes of effectors based on their differential expression in the two types of hosts may suggest varying strategies the pathogen uses to interact with hosts of different susceptibilities. For instance, Class-I effectors might be particularly relevant for the pathogen’s virulence in susceptible hosts, while Class-II effectors might be important for its survival or even virulence in resistant hosts. The significant number of effectors that fall into Class-III suggests the presence of a common set of pathogen strategies that are employed irrespective of the host’s resistance status. Lastly, our study provides a basis for further investigation, as the specific roles of these effectors in pathogenesis remain to be elucidated. Future research may focus on these proteins’ detailed mechanisms, helping us better understand pathogen-host interactions, devise ways to improve host resistance, and develop more effective treatments or preventive measures against infections caused by the Foc strains.

### Carbohydrate active enzymes

Fungal Carbohydrate-Active enZymes (CAZymes) represent a group of enzymes that play crucial roles in the breakdown, biosynthesis, and modification of carbohydrates and glycoconjugates. These enzymes are important in fungal growth, nutrition, and host-pathogen interactions, thus impacting pathogenesis. Carbohydrate-active enzymes (CAZymes) are classified into five distinct modules according to their functional activity and structural resemblance - glycoside hydrolases (GHs), glycosyltransferases (GTs), polysaccharide lyases (PLs), carbohydrate esterases (CEs), and auxiliary activities (AAs). Moreover, a minor fraction of cell wall degrading enzymes (CWDEs) encompasses a non-catalytic module, termed the carbohydrate-binding module (CBMs). Among the numerous CAZymes, a few classes are particularly significant for pathogenesis. For instance, glycoside hydrolases (GHs) and polysaccharide lyases (PLs) are essential for degrading host tissue polysaccharides and subsequent invasion. On the other hand, glycosyltransferases (GTs) contribute to the biosynthesis of fungal cell wall components and various virulence factors. Additionally, carbohydrate esterases (CEs) aid in the modification of plant and fungal cell wall components, often enhancing fungal virulence.

We utilized the domain-based annotation tool, dbCAN, to predict CAZymes in the assembled genomes of IARI-5175 and F-00845, aiming to identify potential CAZyme candidates that could function as CWDEs and contribute to virulence. This process yielded 467 and 494 CAZymes for IARI-5175 and F-00845, respectively, each subsequently divided into different functional groups (Table 4). The majority of these CAZymes were localized on the core chromosomes, specifically 453 (∼97%) in IARI-5175 and 470 (∼95%) in F-00845. Conversely, merely 14 (∼3%) and 24 (∼5%) CAZymes were found on the lineage-specific genomic regions of IARI-5175 and F-00845, respectively. Interestingly, Chromosome-10 in IARI-5175 and Chromosome-3 in F-00845 harboured the most CAZymes. However, notably IARI-5175 contained 315 CWDEs from the functional groups GHs, PLs, CEs, and CBMs, compared to 295 in F-00845.

**Table 4:** Module Wise categorization of CAZymes present in virulent and avirulent Foc strain.

| CAZymes | Virulent | Avirulent |
| --- | --- | --- |

| Modules | Total classes | Total genes | Total classes | Total genes |
| --- | --- | --- | --- | --- |
| AA | 13 | 88 | 13 | 105 |
| CBM | 4 | 11 | 3 | 4 |
| CE | 8 | 31 | 9 | 33 |
| GH | 61 | 250 | 59 | 246 |
| PL | 7 | 23 | 5 | 12 |
| GT | 22 | 64 | 29 | 94 |
| Total | 115 | 467 | 118 | 494 |

Multiple reports have identified CAZymes with “hybrid” functions (Roy et al., 2020; Roy et al., 2021). In this study, we sought to investigate the prevalence of hybrid CAZymes in the genomes of two strains, IARI-5175 (virulent) and F-00845 (avirulent) (Table 5). The term “hybrid” is used to describe CAZymes capable of exhibiting more than one enzymatic activity. This multifunctionality is perceived to offer an evolutionary advantage by enabling an organism to execute several tasks with a single protein. Our investigation led us to identify a distinct presence of hybrid CAZymes in both strains. In total, 8 unique hybrid enzymes were detected in the IARI-5175 genome, while 24 were exclusive to the F-00845 genome. In addition, we noted the presence of 3 hybrid CAZymes shared between both strains. To validate these findings and preclude the possibility that the detected hybrid CAZymes were artifacts of genome assembly, we performed a BLASTP analysis, comparing these hybrid CAZymes with proteins from Fol 4287, a standard reference strain for fungal pathogens. The results revealed a high percentage identity between the hybrid enzymes from our strains and their Fol 4287 counterparts, thus reinforcing the notion that the occurrence of hybrid CAZymes is a common trait among fungal pathogens. Detailed information about these hybrid CAZymes, including their specific chromosome location, the types of catalytic modules they harbor, and the percentage of identity they share with Fol 4287, can be found in Table 5. The identification of these hybrid CAZymes in both fungal strains underlines their potential significance in fungal pathogenesis and points towards the necessity for further research, and perhaps even the development of strategic interventions targeting these enzymes.

**Table 5:**
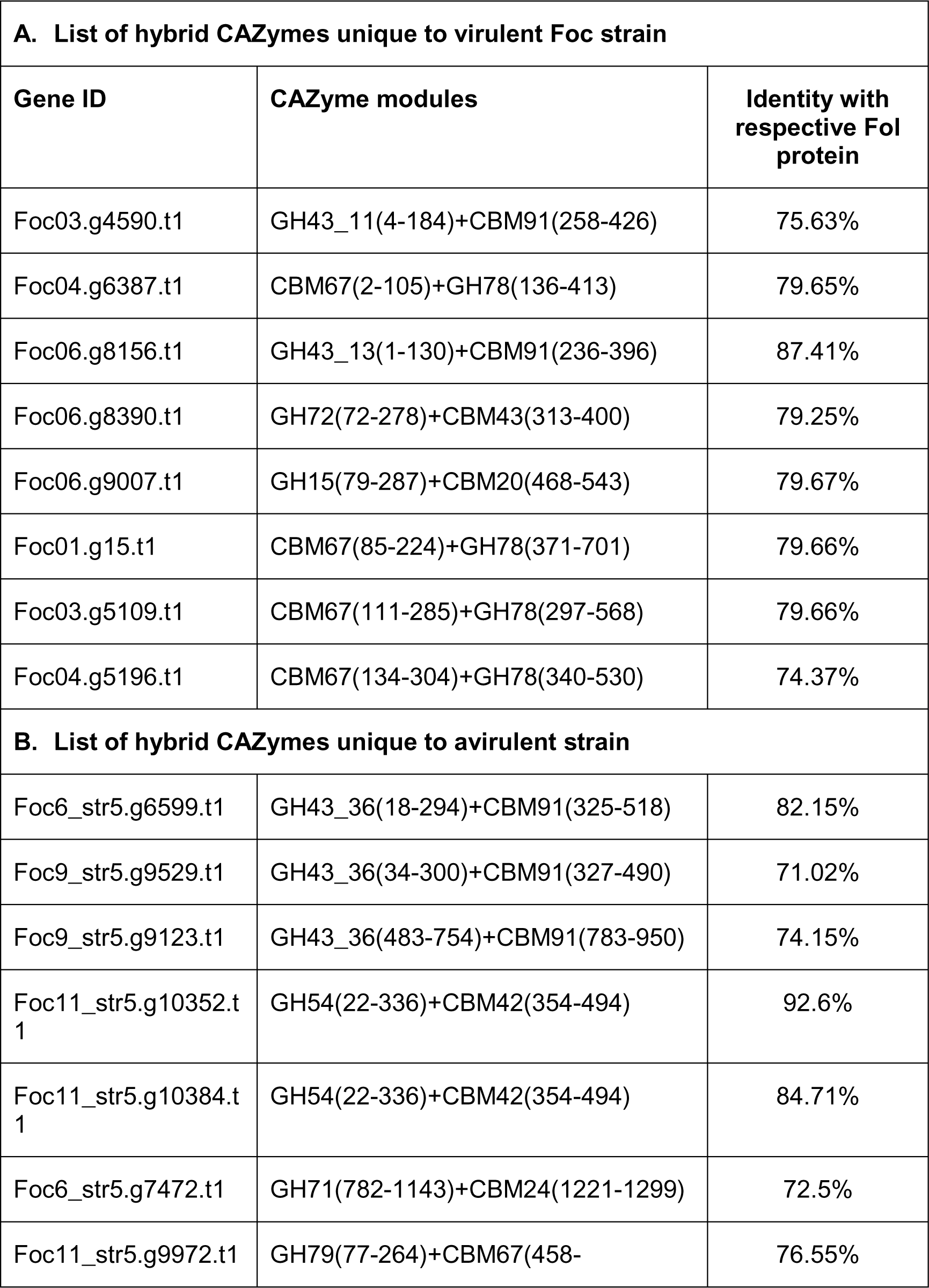
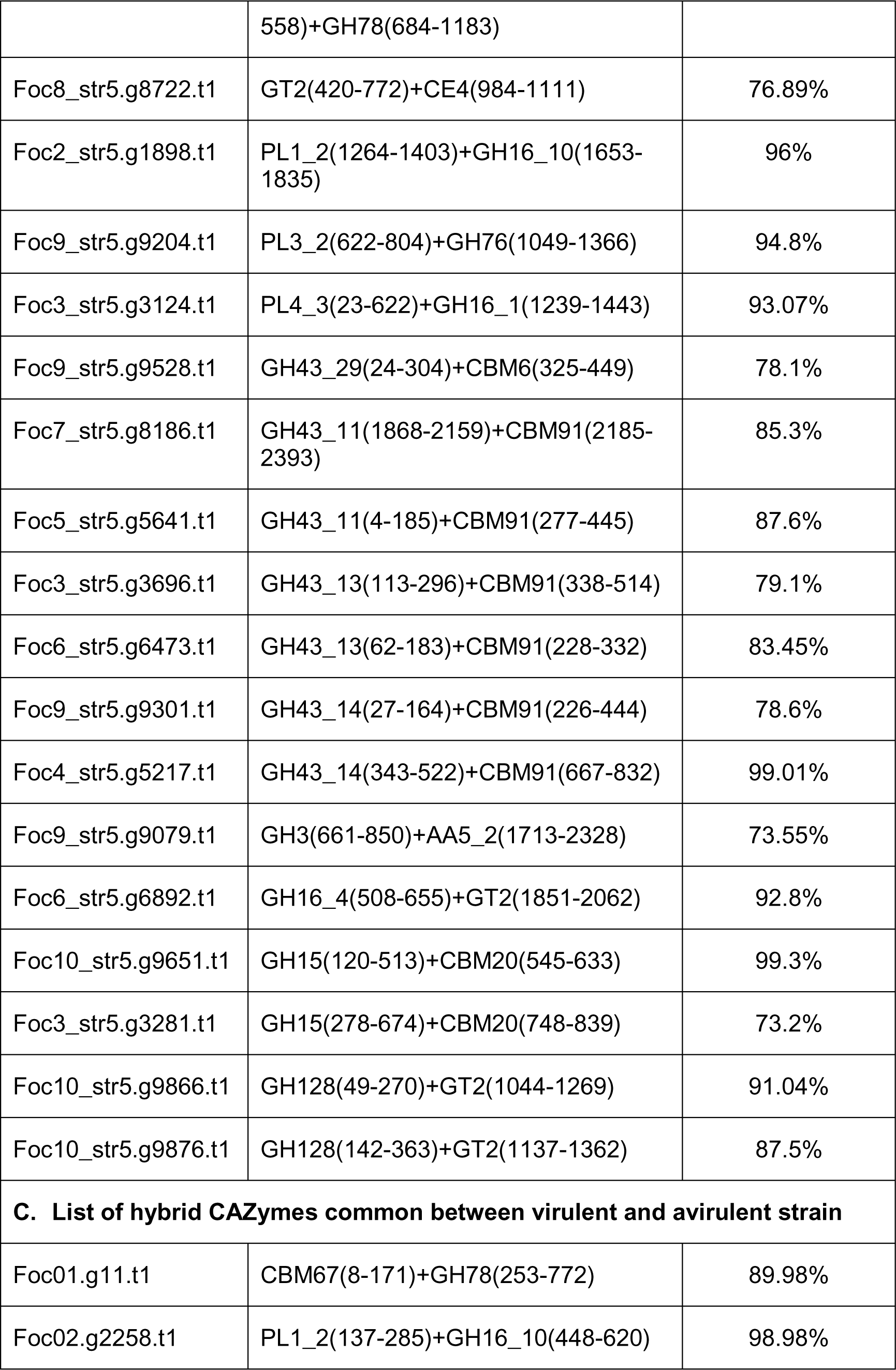
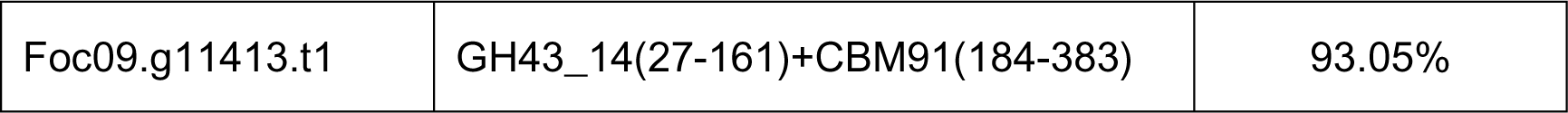
List of hybrid CAZymes predicted in genomes of both virulent and avirulent Foc strains. The gene IDs, CAZymes modules and their % identity with respective Fol protein are listed in column 1, 2 and 3 respectively. A. List of CAZymes unique to virulent strain, B. CAZymes unique to avirulent Foc strain, C. List of CAZymes common between virulent and avirulent Foc strains

The findings from the dbCAN analysis of the genomes of strains IARI-5175 and F-00845 deepen our understanding of the functional characteristics of these organisms, particularly in relation to their potential virulence. The identification of 467 and 494 carbohydrate-active enzymes (CAZymes) in IARI-5175 and F-00845, respectively, underscores the significant roles of these proteins in the organisms’ biology. CAZymes are crucial players in the degradation, modification, and synthesis of carbohydrates, and their roles could extend to virulence, particularly when acting as cell wall-degrading enzymes (CWDEs). It’s noteworthy that more CWDEs were found in IARI-5175 than in F-00845, which could suggest a higher potential for cell wall degradation and hence possibly more efficient infection processes in hosts. The localization of most CAZymes on the core chromosomes of both strains suggests that these functions are central to the organisms’ survival and fitness.

The finding that 301 CAZymes are unique to the IARI-5175 strain could potentially provide insights into this strain’s virulence factors. Among these, the presence of different types of CWDEs exclusive to IARI-5175 reinforces the hypothesis of this strain having an enhanced capability to degrade host cell walls, thus facilitating infection. The identification of certain CAZymes that were exclusively present in the virulent IARI-5175 strain, but absent in the avirulent F-00845 strain, suggests that these proteins might contribute significantly to the pathogenicity of IARI-5175. Further functional analysis and experimental validation of these CAZymes could offer insights into the specific molecular mechanisms behind IARI-5175’s virulence.

On the other hand, the avirulent F-00845 strain also harboured a considerable number of specific CAZymes. The unique CAZymes in F-00845, particularly those related to glycosyltransferases (GTs) absent in IARI-5175, could be interesting targets for further exploration to understand their roles in the avirulence of F-00845. Of particular interest in our research is the discovery of CAZymes with “hybrid” functions. These multifunctional enzymes offer an evolutionary advantage by enabling an organism to perform several tasks with a single protein. In summary, these findings offer valuable insights into the distinct sets of CAZymes in these two strains, opening up new avenues for exploring the mechanisms of virulence and avirulence, and potentially aiding in the development of new strategies for disease control.

To pinpoint CAZymes unique to the virulent IARI-5175 strain, we compared the CAZyme-encoding genes between the two strains. While IARI-5175 shared 166 CAZymes with the F-00845 strain, a substantial tally of 301 CAZymes were exclusive to the IARI-5175 strain. Among the CWDEs, 162 GHs, 20 CEs, 21 PLs, 37 AAs, and 9 CBMs were only present in the IARI-5175 strain. A visual representation of the common and specific CAZymes distribution across chromosomes is shown in Sypplementary Table 3 and Figure 6C. Interestingly, we found that certain CAZymes, including two from the AA class (AA6 and AA12), one each from the CE (CE12) and CBM (CBM91) classes, twelve from the GH family (including GH29, GH51, GH62, GH106, GH114, GH115, GH112, GH131, GH132, GH142, GH162, GH172), and two from the PL class (PL11 and PL 42), were exclusively present in the virulent IARI-5175 strain but absent in the avirulent F-00845 strain. In contrast, no unique GT class CAZymes were identified in the IARI-5175 strain. The avirulent F-00845 strain, on the other hand, had 328 specific CAZymes, comprising 70 AAs, 158 GHs, 22 CEs, 10 PLs, 2 CBMs, and 66 GTs.

**Figure 6:**
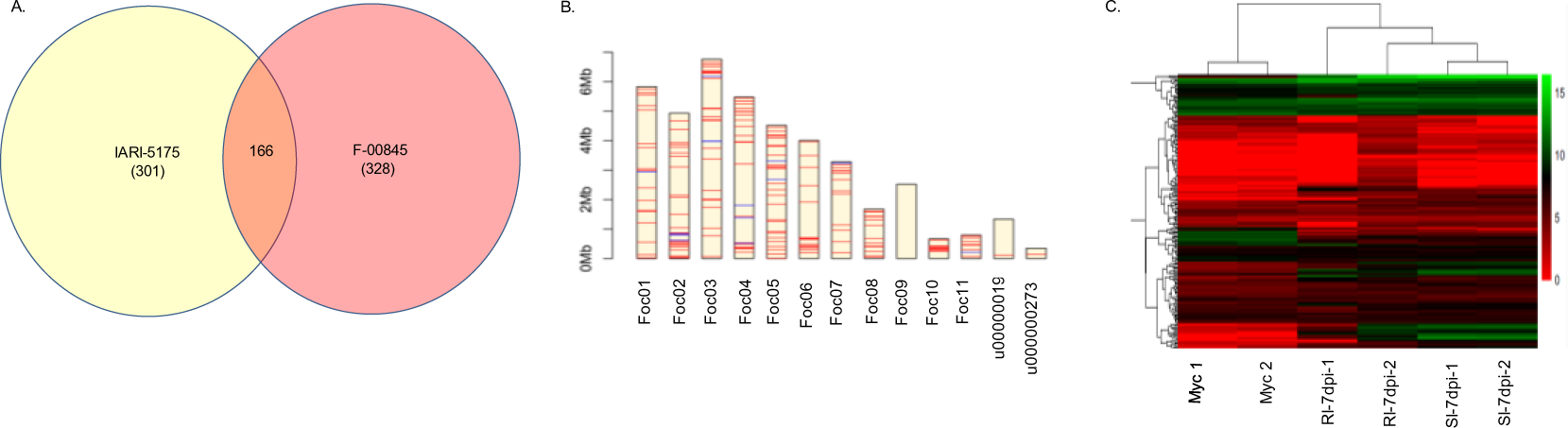
Prediction of CAZymes and their characterization. Genes coding for carbohydrate active enzymes were predicted by dnCAN software, A) Venn diagram of predicted genes coding for CAZymes in IARI-5175 and F-00845 strains. B) Clustering and heatmap of CAZymes present uniquely in IARI-5175 showing expression of effectors in control mycelia, 7 day post infection (dpi) susceptible infected and 7 dpi resistant infected safflower accessions (2 replicates for each treatment). C) Chromosome wise gene block arrangement of CAZymes in IARI-5175.

The comparison of CAZyme-encoding genes between the virulent IARI-5175 strain and the avirulent F-00845 strain is significant in uncovering the potential determinants of virulence. With 301 CAZymes exclusive to IARI-5175, including various CWDEs, it’s clear that this strain possesses a unique set of enzymes that could be involved in its pathogenicity. The existence of certain CAZymes, notably from the AA, CE, CBM, GH, and PL classes, solely in IARI-5175 may suggest that these enzymes could play a crucial role in the strain’s virulence mechanisms. This is consistent with the roles of these classes of CAZymes, which include auxiliary activities (AA) in lignin breakdown, carbohydrate esterases (CE) and glycoside hydrolases (GH) involved in polysaccharide degradation, and polysaccharide lyases (PL) and carbohydrate-binding modules (CBM) associated with substrate binding and specificity. Interestingly, the virulent IARI-5175 strain doesn’t possess any unique GT class CAZymes, which are primarily involved in the biosynthesis of complex carbohydrates and glycoconjugates. This absence may suggest that unique GTs are not crucial for the virulence of IARI-5175, or that common GTs shared with F-00845 may be sufficient for its needs. On the contrary, the avirulent F-00845 strain possesses a substantial number of unique CAZymes, including 66 GTs. The high proportion of unique GTs in F-00845 might suggest their importance in maintaining this strain’s non-pathogenic nature, possibly by focusing on self-growth and survival, rather than host interactions. In conclusion, these findings provide a significant leap towards understanding the CAZymes-mediated virulence of the IARI-5175 strain and avirulence of the F-00845 strain. Unravelling these differences could lead to the identification of potential targets for mitigating the pathogenicity of virulent strains like IARI-5175. Furthermore, it may help in the development of novel strategies for disease management and prevention by capitalizing on our knowledge of relative differences in the expression of CAZymes between the pathogenic and non-pathogenic strains.

Next, we evaluated the expression of the 166 common and 301 IARI-5175 strain-specific CAZymes in the transcriptome data. Out of the 301 unique CAZymes, 118 exhibited differential regulation in compatible disease interactions with safflower. These included, a total of 95 CAZyme-encoding genes spanning all 6 classes (AA, CE, CBM, GH, GT, PL) which showed upregulation, and 23 CAZyme-encoding genes belonging to 4 CAZyme classes (AA, CE, GH, GT) which displayed down regulation. The majority of upregulated differentially expressed genes (DEGs) during compatible interaction were from the CAZyme family GHs (63), followed by AAs (12), PLs (12), and CEs (5). The fewest upregulated DEGs were observed in classes GT (2) and CBM (1). Among the down regulated genes, the GH class had the highest number (14), followed by AA and GT with 4 genes in each category, and CE (1).

During incompatible disease interaction, DEGs from only four classes (GH, PL, CE, AA) showed upregulation, while three classes (GH, GT, AA) demonstrated down regulation. Specifically, 75 CAZymes exhibiting differential gene expression were identified in the resistant interaction of Foc with Sharda. In total, 63 CAZymes, encompassing GHs (45), PLs (8), CEs (5), and AAs (5), showed upregulated gene expression during the resistant interaction. Conversely, 12 CAZyme-encoding genes, belonging to GHs (8), GTs (2), and AAs (2), were down regulated during the same interaction.

In the 166 Carbohydrate-Active Enzymes (CAZymes) common to both virulent and avirulent strains, differential expression was observed in 99 and 54 CAZyme-encoding genes during compatible and incompatible interactions, respectively. In the compatible interaction, 68 CAZymes that showed upregulated gene expression were primarily from GH (40) and AA (13) classes followed by CE (7), GT (6), CBM (1), and Pl (1). Conversely, 31 CAZyme-encoding genes from GH (16), GT (8), and AA (7) classes were down regulated. During the incompatible interaction, 39 CAZyme-encoding genes from the GH (23), CE (6), AA (5), GT (2), PL (1), and CBM (1) classes exhibited upregulation, while 15 CAZyme-encoding genes from the GH (8), GT (4), and AA (3) classes were down regulated.

Interestingly, 44 CAZymes [AA (5), CBM (1), CE (4), GH (26), GT (2), PL (6)] were not expressed in the fungal mycelia but showed remarkable upregulation during compatible interaction. Of these 44 CAZymes, 43 were differentially regulated during the incompatible interaction. Three genes [AA (1) and GH (2)] displayed a negative correlation in their expression, being down regulated in the incompatible interaction but upregulated in the compatible interaction. The remaining 40 CAZymes showed a positive correlation, with upregulation in both types of interactions. One specific CAZyme (gene ID Foc01.g11), with significant upregulation in the compatible interaction, had no expression either in the mycelia or during incompatible interaction. This gene is a hybrid CAZyme with CBM and GH catalytic modules. Despite the identification of 14 CAZyme-encoding genes within the accessory genome of the virulent strain, none of them demonstrated differential expression in either the compatible or incompatible interactions.

Our comprehensive evaluation of the expression of common and strain-specific carbohydrate-active enzymes (CAZymes) in the transcriptome data brings us one step closer to deciphering the mechanisms underlying the virulence of the IARI-5175 strain. Among the 301 unique CAZymes in IARI-5175, 118 showed differential regulation during disease interaction with susceptible safflower, indicating their active role during host invasion. The upregulation of 95 CAZymes, primarily from the glycoside hydrolase (GH) class, underlines their potential role in breaking down host cell wall components, promoting pathogen colonization. On the other hand, the resistant disease interaction revealed differential expression in fewer CAZymes (75). The majority of these belonged to the GH class, which also showed upregulation, indicating a possible defence mechanism of the resistant host in countering the fungal attack. The 166 CAZymes common to both virulent and avirulent strains showed differential expression during both compatible and incompatible interactions.

These shared CAZymes could be playing basic roles required by the fungus during host invasion, irrespective of the host’s resistance status. Interestingly, 44 CAZymes were not expressed in the fungal mycelia but showed significant upregulation during compatible interaction. This indicates that these CAZymes might be crucial during the initial stages of infection and could be potential targets for controlling the disease. The presence of some genes showing opposite expression patterns between the compatible and incompatible interactions may suggest their specific roles in determining the outcome of disease interaction. The identification of a hybrid CAZyme (Foc01.g11) with significant upregulation in the compatible interaction but no expression during incompatible interaction or in the mycelia is intriguing. This CAZyme may be a crucial player in the virulence of the IARI-5175 strain, requiring further investigation to understand its specific function. Interestingly, none of the 14 CAZyme-encoding genes identified within the accessory genome of the virulent strain demonstrated differential expression in either types of disease interaction. This observation suggests that the virulence of the IARI-5175 strain may not be solely dependent on these accessory CAZymes. Overall, these findings shed light on the complex interplay between the fungal pathogen and its host at the molecular level, revealing a dynamic regulation of CAZymes that appears to be tailored to the host’s resistance status.

### Secondary metabolites

Fungal secondary metabolite gene clusters play a critical role in plant pathogenesis. These clusters encode a suite of secondary metabolites that act as virulence factors, promoting infection and disease development in host plants. Such metabolites can interfere with plant defence responses, modify plant tissues to facilitate fungal colonization, and directly damage plant cells. A notable example is the biosynthesis of host-selective toxins by certain plant-pathogenic fungi, which are toxic only to susceptible cultivars of the host plant and are crucial for disease development. These toxins are produced by specific gene clusters in the fungal genome. Moreover, the expression of these clusters is typically tightly regulated, being induced during the infection process and in response to plant-derived signals. Understanding the regulation, biosynthesis, and function of these secondary metabolite gene clusters can provide valuable insights into the mechanisms of fungal plant pathogenesis and can contribute to the development of effective disease management strategies. Secondary metabolite gene clusters in the genome of strain IARI-5175 and F-00845 were searched using antiSMASH software. A total of 28 and 45 gene clusters were identified in the virulent strain Foc-IARI-5175 and the avirulent strain F-00845, respectively. These clusters were categorized into six different classes (Table 6). Both strains exhibited the highest number of Non-ribosomal peptide synthetase (NRPS) clusters, followed by Polyketide synthase (PKS), terpenes, hybrid, and Ribosomally synthesized and post-translationally modified peptide products (RiPP). In strain IARI-5175, all 28 gene clusters were located on the core genome, while in strain F-00845, secondary metabolite gene clusters were distributed on both the core (42) and accessory (3) genomes. Gene clusters for fusaric acid, a characteristic fungal mycotoxin of the Fusarium group, were found on chromosome-4 in both Foc strains IARI-5175 and F-00845. However, the Foc strains differed in their mycotoxins, with beauvericin found in IARI-5175 (Chromosome-8) and ACT-toxin found in F-00845 (Chromosome-10). A comparative chromosome-wise distribution of secondary metabolite genes for IARI-5175 and F-00845 is shown in Figure 7. A comparison was made between genes coding for secondary metabolites in IARI-5175 and F-00845 to identify unique and shared genes in the virulent strain IARI-5175. The virulent strain contained 222 unique secondary metabolite-encoding genes and shared 156 genes with F-00845. Conversely, 375 genes coding for secondary metabolites were unique to the F-00845 strain.

**Figure 7:**
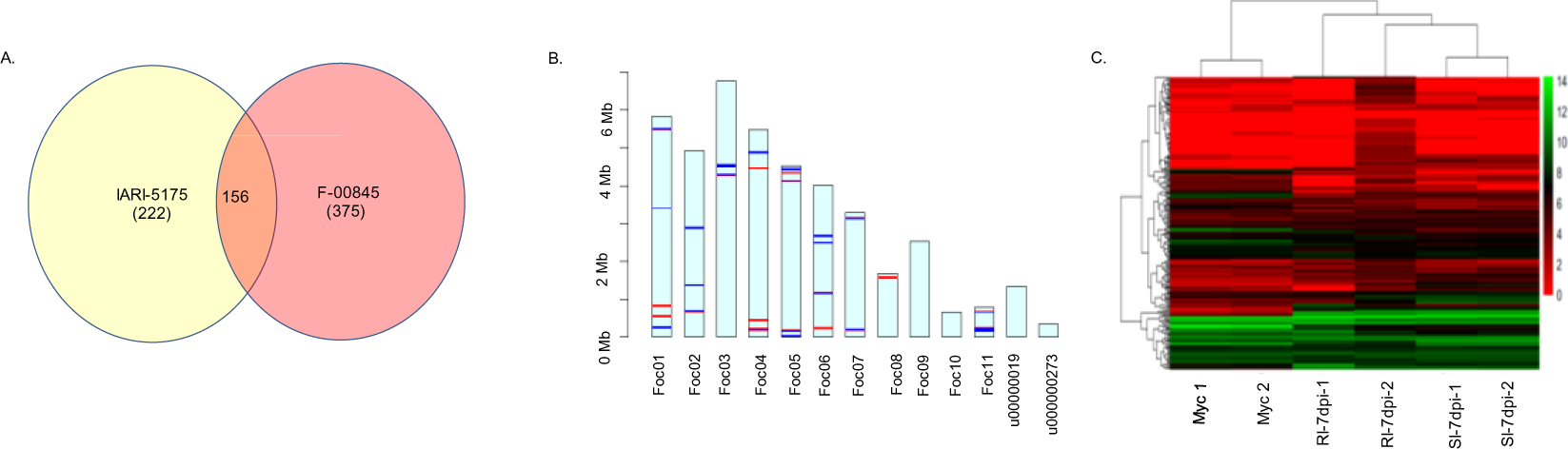
Prediction of Secondary metabolite gene clusters and their characterization. Genes coding secondary metabolites were predicted by antiSMASH software, A) Venn diagram of predicted genes coding for secondary metabolite in IARI-5175 and F-00845 strains. B) Clustering and heatmap of secondary metabolites present uniquely in IARI-5175 showing expression of effectors in control mycelia, 7day post infection (dpi) susceptible infected and 7 dpi resistant infected safflower accessions (2 replicates for each treatment). C) Chromosome wise gene block arrangement of secondary metabolite gene clusters in IARI-5175.

**Table 6:** Distribution of the secondary metabolite clusters in virulent (V) and avirulent (AV) strains of Foc.

| S. No. | Secondary metabolite cluster | V |  | AV |  |
| --- | --- | --- | --- | --- | --- |
|  |  | Core | LS | Core | LS |
| 1 | Non ribosomal peptide synthetase (NRPS) | 11 | 0 | 14 | 2 |
| 2 | Ribosomally synthesised and post-translationally modified peptide product (RiPP) | 2 | 0 | 4 | 0 |
| 3 | Terpes | 5 | 0 | 8 | 0 |
| 4 | Polyketide synthase (PKS) | 7 | 0 | 9 | 0 |
| 5 | Hybrid | 3 | 0 | 6 | 1 |
| 6 | Other | 0 | 0 | 1 | 0 |

We further leveraged RNAseq data to scrutinize the expression of genes linked to the synthesis of both unique and shared secondary metabolites during compatible and incompatible interactions. Out of the 156 shared secondary metabolite genes evaluated, 99 genes were found differentially expressed during compatible interactions, with 51 upregulated and 48 down regulated. However, during incompatible interactions, 45 genes displayed differential expression, with 23 upregulated and 22 down regulated. On inspecting the expression of secondary metabolite genes exclusive to the IARI-5175 strain, it was discerned that 72 genes in the compatible interaction and 39 in the incompatible interaction manifested differential expression. When Foc interacted with the susceptible host, there was an upregulation of 39 genes and down regulation of 33. Simultaneously, the interaction between Foc and the resistant host resulted in upregulation and down regulation of 23 and 16 secondary metabolite genes respectively.

The secondary metabolite gene clusters connected with the mycotoxins beauvericin and fusaric acid, comprising 21 and 19 genes respectively, drew particular interest. In the susceptible host infected by Foc, 12 genes related to beauvericin showed differential expression, with 7 down regulated and 5 upregulated. However, only 7 DEGs related to beauvericin were found to be regulated during Foc’s infection with the resistant host, with 4 upregulated and 3 down regulated genes. Among the 19 genes associated with fusaric acid synthesis and transport, 4 were down regulated and 1 upregulated in the compatible interaction, but no DEGs were noticed in the resistant host during the incompatible interaction with safflower. A gene, Foc04.g5197, upregulated during the compatible interaction and coding for fusaric acid, didn’t express in either control mycelial tissue or during the incompatible interaction. Furthermore, Foc04.g5197 was functionally annotated as a putative transporter responsible for the export of fusaric acid out of the cell. This suggests the potential significance of this gene in the transport of fusaric acid and its possible contribution to Foc’s virulence against safflower.

Within the beauvericin biosynthetic gene cluster, two genes displayed an inverse correlation in compatible and incompatible interactions. These genes were down regulated in the compatible interaction and showed no differential regulation in the incompatible interaction. A functional annotation via BLAST search disclosed that one of these down regulated genes codes for Beauvericin nonribosomal cyclodepsipeptide synthetase, crucial for the rapid synthesis of beauvericin through a nonribosomal, thiol-templated mechanism. The other down regulated gene coded for a lysyl oxidase-like protein, which facilitates the essential cross-linking formation of the extracellular matrix (ECM) through catalysing the oxidative deamination of lysine side chains on collagen and elastin. Contrastingly, two genes upregulated during compatible interactions didn’t display differential expression in incompatible interactions. Functional annotation of these genes determined that one coded for a protein with an unknown function, while the other coded for beauvericin cluster specific repressor BEA4. This suggests that beauvericin biosynthesis might be suppressed during compatible interactions at later stages of Fusarium wilt disease.

Our work holds profound significance in a number of ways, as it delves deeply into the mechanisms of plant-fungal interactions and the role of secondary metabolites in pathogenesis. At the heart of the findings is the substantial role fungal secondary metabolites play in fostering infection and disease progression in host plants. The study underscores the destructive power of these metabolites, whether it’s modifying plant tissues to ease fungal colonization, hindering plant defence mechanisms, or directly assaulting plant cells. The research also makes significant strides in the study of secondary metabolite gene clusters in fungi. We found these clusters to be central to the production of host-selective toxins, key contributors to disease progression. This expression of these clusters is a meticulous process, often induced during infection and in response to plant-derived signals. This discovery could yield a plethora of information on the orchestration of plant-pathogen interactions. But the significance of our study doesn’t stop with an understanding of these processes, it also provides a stepping stone to the development of disease management strategies. The knowledge gained on the functions and regulation of these gene clusters can be exploited to devise novel disease management strategies, making this study not only insightful but also practical.

Moreover, the comparative analysis of two fungal strains, the virulent strain IARI-5175 and the avirulent strain F-00845, has brought to light key differences in secondary metabolite gene clusters. The RNAseq data analysis further provides a nuanced understanding of gene expression patterns during compatible (leading to disease) and incompatible (leading to resistance) plant-pathogen interactions. It’s these differential patterns in the expression of secondary metabolite genes that could be potentially manipulated to develop durable resistance in crops against devastating diseases. Particularly intriguing is the discovery of the role of the gene Foc04.g5197 in the transport of fusaric acid, a potent mycotoxin. The observed upregulation of this gene during infection of the susceptible host suggests its possible contribution to the virulence of Fusarium. This could open new avenues in targeted gene manipulation for disease management.

Additionally, the work conducted on the beauvericin biosynthetic gene cluster gives an insight into the complex regulation of fungal toxins and presents potential targets for developing disease resistance. The contrasting regulation of certain genes in this cluster during compatible and incompatible interactions further emphasizes the dynamic nature of host-pathogen interactions. In conclusion, the importance of our study lies in the comprehensive picture it paints of the intricate and tightly regulated fungal secondary metabolite gene clusters. It emphasizes the roles these clusters play in fungal virulence and disease development, and lays the foundation for the future development of effective disease management strategies.

#### 3.3.9 Comparative transcriptome analysis during resistant and susceptible interaction

Our research aimed to identify Foc genes with expression patterns distinctively influenced by susceptible and tolerant hosts. We performed transcriptome sequencing on Foc mycelia from the IARI-5175 strain, as well as on infected root tissues from hosts exhibiting contrasted reactions to infection. High Pearson’s Coefficients of 0.97, 0.98, and 0.89 for mycelia, infected susceptible host (compatible interaction), and infected tolerant host (incompatible interaction) respectively, suggested a strong similarity between the biological replicates. The data was analysed to identify 3 groups of genes: Group A (Mycelia vs Compatible Interaction), Group B (Mycelia vs. Incompatible Interaction), and Group C [Differentially expressed genes (DEGs) of Group A vs DEGs of Group B]. Group A analysis revealed 5694 Foc genes demonstrating differential regulation in response to infection in the susceptible host (PI-199897). These encompassed 3026 up regulated and 2668 down regulated genes (Figure 3.9A). Of the up regulated genes, 260 were exclusive to infection, showing no expression in mycelial tissue. The remaining 2766 up regulated genes exhibited expression in both infection and mycelial contexts. Comparably, among the down regulated genes, 80 “localization”, “establishment of localization”, and “transport”. We noted that many of these genes were linked to the transport of various organic molecules and ions in the susceptible host. Likewise, the molecular functions associated with upregulated genes revolved around transporter activities, encompassing transmembrane transporter activity and symporter activity (Figure 8B). Similarly, the down regulated genes were associated with the GO terms for biological processes (508 genes), cellular components (281 genes), and molecular functions (1841 genes). The highest number of these genes were related to biological GO terms like “small molecule metabolic processes”, “mitotic cell cycle metabolic processes”, and “DNA metabolic processes”, with 87, 75, and 54 genes respectively in each category. Additional enrichment was observed for “DNA replication initiation” and “DNA repair”, with 13 and 41 genes, respectively. The molecular functions linked with down regulated genes encompassed “ion binding”, “protein binding”, and “small molecule binding”, with counts of 195, 185, and 130 genes respectively (Figure 8C). Group B analysis revealed that in the resistant host infected by Foc, there was a lower count of differentially expressed genes (2210) compared to the susceptible host. From these, 1350 genes were up regulated and 860 were down regulated (Figure 9A). Among the up regulated genes, the expression of 163 was uniquely induced upon infection, as they were not expressed in the mycelial tissues. The rest, amounting to 1187 genes, were present in mycelia and saw a significant increase in expression during the incompatible interaction. On the other hand, among the down-regulated genes, only five showed expressions exclusive to the mycelia. The expression of the remaining 855 genes decreased upon infection compared to their mycelial expression.

**Figure 8:**
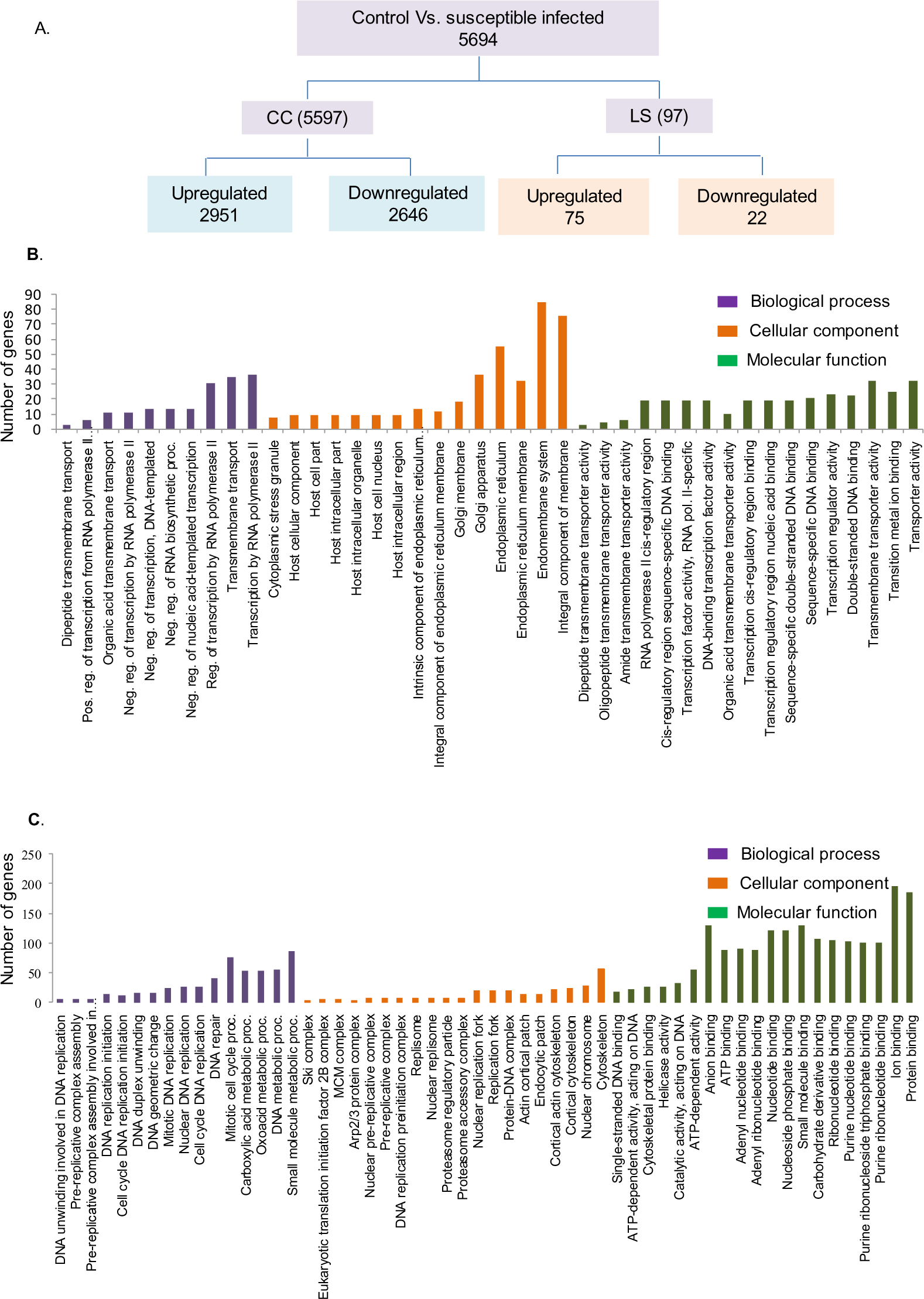
Differentially regulated genes during control Vs susceptible infected safflower. A) Detail of DEGs in core and lineage specific genome. B) GO enrichment analysis of upregulated in DEGs. C) GO enrichment analysis of downregulated in DEGs.

**Figure 9:**
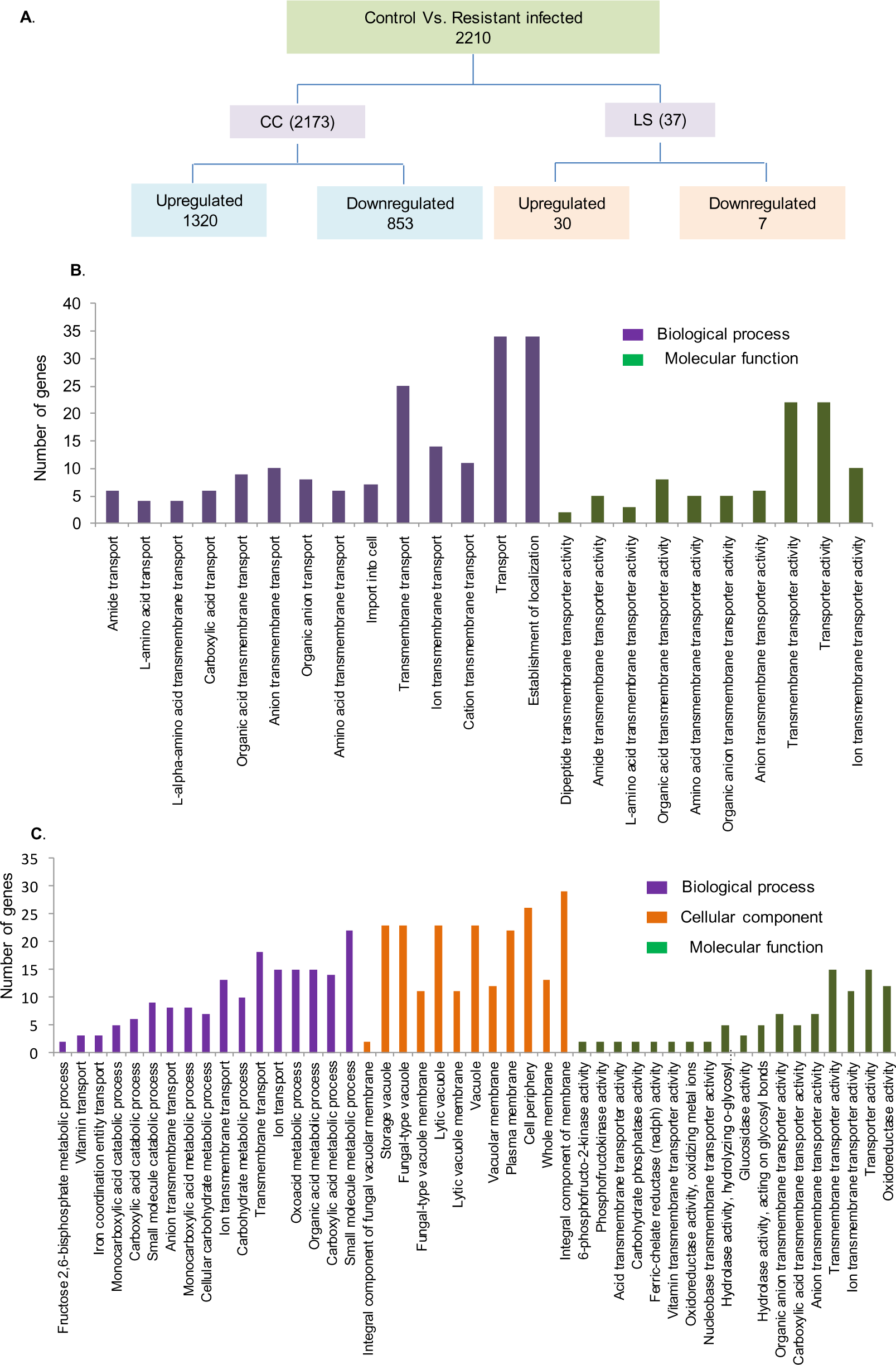
Differentially regulated genes during control Vs Resistant infected safflower. A. Detail of DEGs in core and lineage specific genome. B. GO enrichment analysis of upregulated in DEGs. C. GO enrichment analysis of downregulated in DEGs.

GO enrichment analysis of upregulated DEGs in the resistant host revealed that these genes were primarily associated with biological functions akin to those observed in the susceptible host. These functions were categorized by GO terms such as “transport”, “establishment of localization”, and “transmembrane transport”, with 32, 32, and 24 genes in each category, respectively. Additionally, 135 upregulated genes were related to diverse molecular pathways, mostly associated with transmembrane transporter activity of various molecules, including ions, inorganic entities, and anions (Figure 9B). Contrarily, down regulated genes in the resistant host displayed enrichment in GO terms like “small molecule metabolic process” and “transmembrane transport”, with 22 and 18 DEGs in each category, respectively. These genes were majorly associated with categories like molecular transport (including vitamins, iron coordination entities, ions), metabolic processes (e.g., fructose 2,6-bisphosphate, monocarboxylic acid, cellular carbohydrate, organic acid, carboxylic acid, and small molecules), and catabolic processes (involving monocarboxylic acid, carboxylic acid, and small molecules). From the perspective of molecular pathways, the most substantial number of genes were linked with GO terms like “transmembrane transporter activity”, “oxidoreductase activity”, and “hydrolase activity acting on glycosyl bonds”. Additional molecular pathways, such as “6-Phosphofructo-2-Kinase Activity” and “phosphofructokinase activity”, also showed enrichment with two genes for each GO term (Figure 9C).

To determine differentially expressed genes (DEGs) with either similar or inversely correlated expression patterns during compatible (susceptible host) and incompatible (resistant host) interactions, the DEGs from each host type were compared (Group C comparison). From the 5694 DEGs observed in the compatible interaction and the 2210 from the incompatible interaction, 1970 common genes were found to be differentially regulated in both interactions. A positive correlation was seen in the expression of 995 of these genes, while 975 displayed an inverse correlation between the two interactions. Moreover, out of the 995 positively correlated genes, 596 were up regulated and 399 were down regulated in both interactions. Interestingly, among the inversely correlated genes, 561 were up regulated in the compatible host but down regulated in the incompatible host, and 414 showed the opposite pattern: down regulation in the compatible host and up regulation in the incompatible host. GO terms for all DEGs are listed in “Supplementary Table 6”.

GO enrichment analysis revealed that the majority of the commonly up regulated genes were grouped under GO terms for molecular pathways such as “catalytic activity”, “lyase activity”, and “binding”, each having 15, 7, and 7 genes respectively. Furthermore, these up regulated genes were predominantly localized in “cellular anatomical entity” (13 genes), extracellular region (8 genes), and host cellular region (12 genes). The key biological functions associated with these up regulated genes were metabolic processes, catabolic processes, cell wall organization, and external encapsulating structure organization) (Figure 10A).

**Figure 10:**
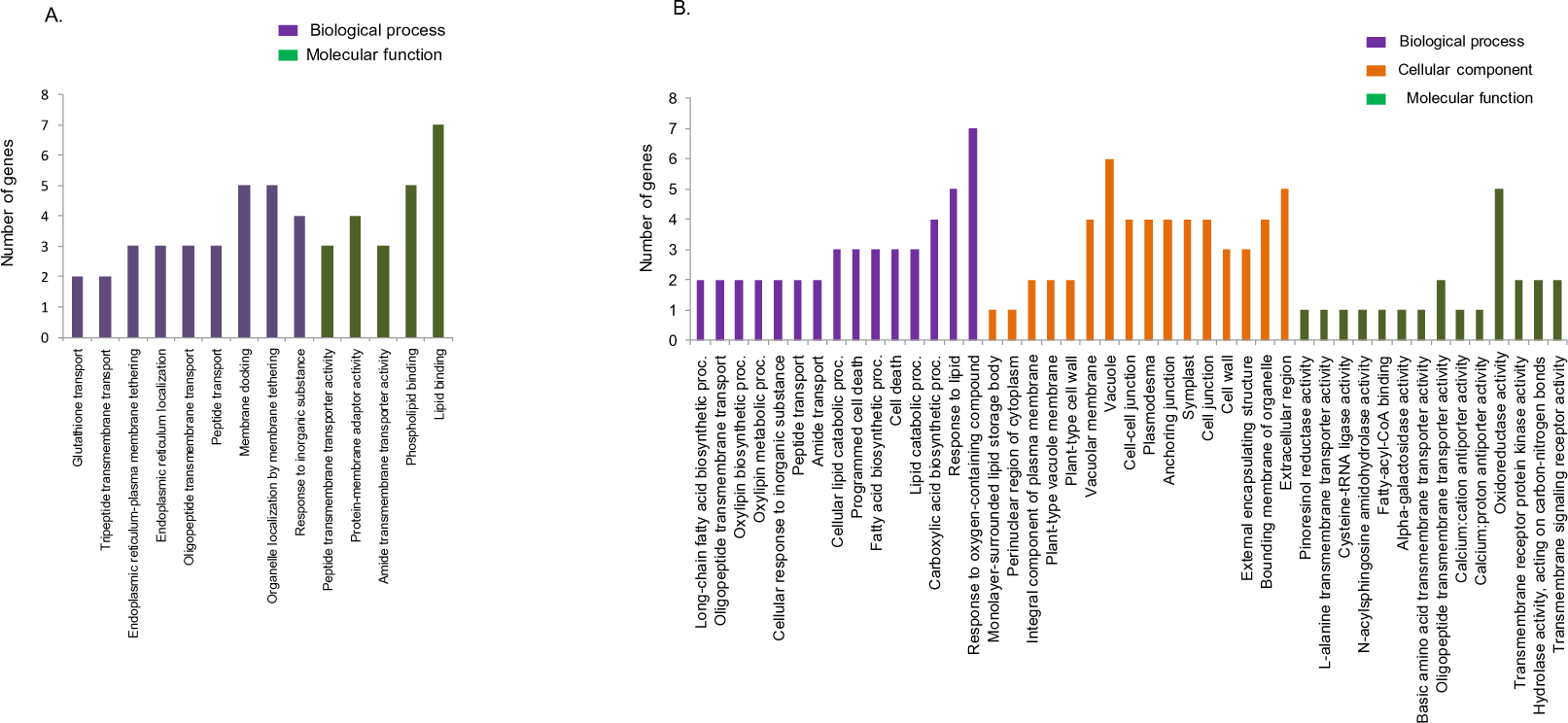
GOs of the positively DEGs from compatible and incompatible interactions. A) GO enrichment analysis of DEGs commonly upregulated in SI and RI. B) GO enrichment analysis of DEGs commonly down regulated in SI and RI

Down regulated genes were primarily grouped under GO terms for molecular functions like “lipid binding” (7 genes), “phospholipid binding” (5 genes), “protein-membrane adaptor activity” (4 genes), “peptide transmembrane transporter activity” (3 genes), and “amide transmembrane transporter activity” (3 genes). Intriguingly, no cellular component demonstrated enrichment for down regulated genes in either the susceptible infected or resistant infected host. However, biological functions tied to 30 up regulated genes were grouped under GO terms such as “membrane docking” (5 genes), “organelle localization by membrane tethering” (5 genes), and “response to inorganic substance” (4 genes). The GO terms associated with the commonly upregulated and down regulated DEGs are presented in Figure 10A and 10B respectively.

Of the genes that were up regulated in the compatible host but down regulated in the incompatible host, the most were associated with GO terms “transporter activity” (13 genes) and “transmembrane transporter activity” (12 genes). These DEGs were primarily found in the intrinsic component of the membrane (23 genes) and integral component of the membrane (22 genes). The biological processes associated with these genes included localization, transport, establishment of localization, and transmembrane transport. Conversely, for genes that were down regulated in the compatible host and up regulated in the incompatible host, the majority were categorized under GO terms related to molecular pathways such as “nucleotide binding” (25 genes), “nucleoside phosphate binding” (25 genes), and “adenyl ribonucleotide binding” (18 genes). The associated biological processes were primarily organized under GO terms “mitotic cell cycle process” and “mitotic cell cycle”, each with 17 genes. The GO terms associated with the negatively regulated DEGs are presented in Figure 11 A and 11 B respectively.

**Figure 11:**
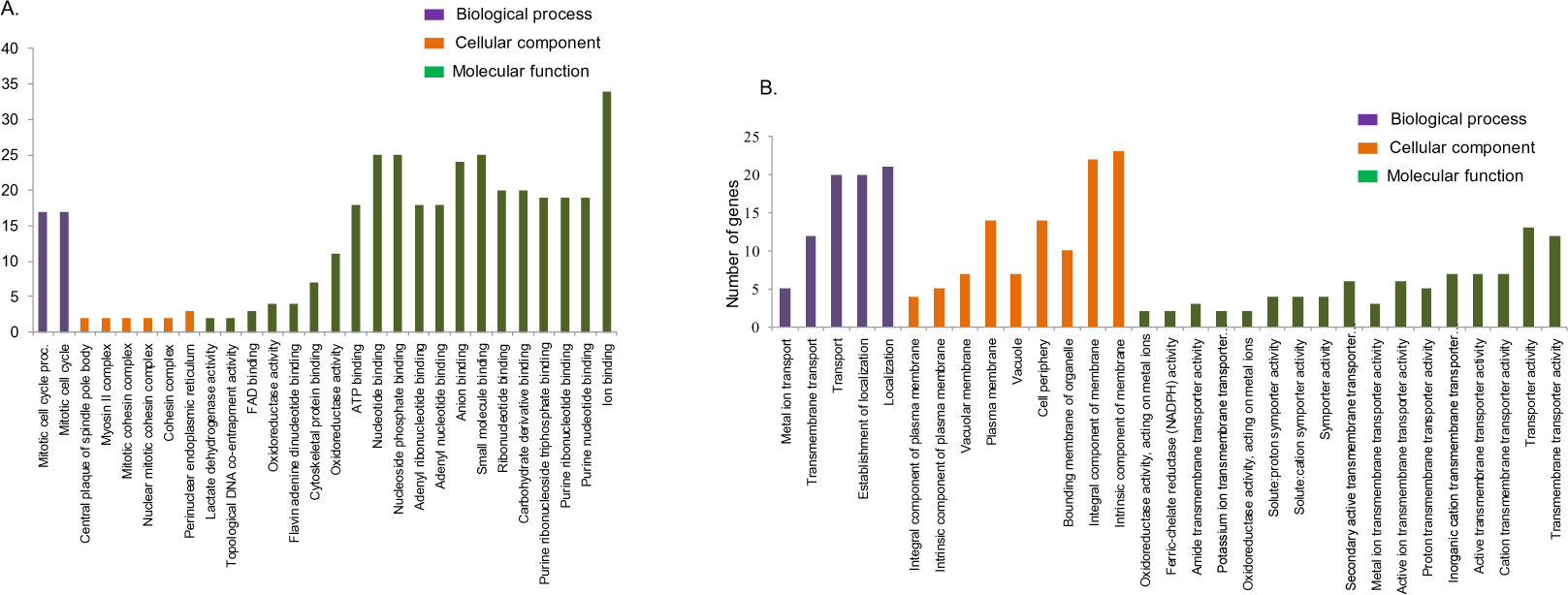
GOs of the inversely correlated DEGs from compatible and incompatible interactions. A) GO enrichment analysis of DEGs upregulated in SI and downregulated in RI. B) GO enrichment analysis of DEGs down regulated in SI and upregulated in RI.

Our study provides crucial insights into the gene expression changes occurring in the Foc fungus during infection of susceptible and tolerant hosts. The transcriptome sequencing of Foc mycelia from the IARI-5175 strain and infected root tissues revealed distinct transcriptional patterns reflective of the disease outcome. The fact that there were more differentially expressed genes (DEGs) in the compatible interaction (susceptible host) compared to the incompatible interaction (resistant host) may indicate that the fungus alters its gene expression more dramatically in the susceptible host, where it can thrive. Interestingly, both up- and down regulated genes were observed in both hosts. Genes showing exclusive up regulation during infection might be related to the pathogenic activity of the fungus, such as host invasion, while those being down regulated could be associated with routine functions not required during infection. Enrichment analyses of these DEGs suggested a few common patterns: “transport” and “localization” were notably up regulated processes, possibly reflecting the fungus’s need to transport nutrients and other essential molecules across the host-fungal interface. Down regulated genes in both host types showed enrichment for processes related to cell metabolism, possibly reflecting a shift from fungal growth and maintenance to disease-promoting activities.

The analysis also revealed some striking differences in the Foc gene expression during the compatible and incompatible interaction. Despite similar GO categories, there were fewer DEGs in the incompatible interaction, indicating a less dramatic transcriptional response, possibly due to the host’s resistance mechanisms. Most intriguing, some genes showed an inverse correlation between the two interactions, being up regulated in the susceptible host and down regulated in the resistant host, and vice versa. This could reflect different strategies used by the fungus in response to host resistance. These inversely correlated genes may have crucial roles in determining the disease outcome, suggesting they might be of interest as targets for disease control. In conclusion, our work provides comprehensive insights into the dynamic transcriptional reprogramming that Foc undergoes during host invasion. This could be instrumental in understanding the molecular mechanisms underlying Foc virulence and developing novel disease management strategies.

## 4 Discussion

In an effort to understand the molecular mechanisms driving the pathogenicity of *F. oxysporum* f. sp. *cartham*i (Foc), a fungal pathogen with significant impacts on safflower crops in India, we employed a comparative approach using genomics and transcriptomics. This analysis focused on the comparison of Foc strains exhibiting stark differences in virulence on safflower plants. Our investigation began with the pathogenicity testing of 16 Foc isolates on two accessions of safflower: one susceptible (PI-199897) and one resistant (Sharda) to Foc. This allowed us to categorize the Foc strains into two groups - virulent and avirulent - with each group composed of 8 Foc strains. We also examined the Foc strains morphologically at both the macroscopic and microscopic levels, focusing on traits such as colony morphology, pigmentation, spore dimensions, sporulation, spore germination, and growth rate. The aim was to determine whether morphological differences corresponded to variations in the pathogenic potential of the Foc strains. Our results showed that one of the virulent Foc strains, F-00940, had the slowest growth rate at 4.53 mm/day, while an avirulent Foc strain, F-00933, displayed the highest growth rate at 6.98 mm/day. We observed variations in other morphological traits as well. However, these differences did not clearly distinguish between avirulent and virulent Foc strains. As such, we found no clear correlation between the different pathotypes (virulent and avirulent) and the morphology of the Foc strains. Previous studies have reported substantial morphological and genetic diversity among *F. oxysporum* strains. For instance, an extensive diversity analysis on Indian Foc isolates was performed by Singh et al (2019). Despite these reports, no confirmed correlation exists between virulence and morphological traits in Foc. Similar studies have also found considerable variability in morphology and virulence in other formae speciales, yet no connection was established between these traits (Dubey et al., 2010). The lack of a clear correlation between virulence and morphology could potentially be due to the influence of a combination of genetic and environmental factors, such as light and pH, on fungal strain morphology, independent of their virulence (Pradeep et al., 2013).

Our investigation continued with a phylogenomic analysis to unravel the evolutionary relationships among Foc strains and closely related formae speciales. The fact that all thirteen Foc strains in our study form a single clade suggests their monophyletic origin. Interestingly, virulent Foc strains didn’t cluster separately from avirulent strains, but rather, they were interspersed. Despite this common ancestry, the scattered placement of virulent and avirulent strains within the Foc clade indicates separate evolutionary trajectories, contributing to varied pathogenicity. This is indicative of the evolutionary plasticity within this species complex, possibly reflecting the organism’s adaptability to different environments, hosts, and pathogenic circumstances. The homogenous virulence phenotype against safflower seems to be a result of convergent evolution, rather than a uniform genomic characteristic. However, our findings warrant further research involving a larger population of Foc strains to provide more definitive evidence supporting the evolution of Foc. It’s noteworthy that previous phylogenetic studies have confirmed *F. oxysporum* f. sp. ciceris as a monophyletic group (Jiménez-Gasco et al., 2002), which is in agreement with its genetic kinship with *F. oxysporum* f. sp. carthami. On the other hand, *F. oxysporum* f. sp. lycopersici and *F. oxysporum* f. sp. vasinfectum were proposed to have polyphyletic origins (Skovgaard et al., 2001). The pathway by which Fusarium strains acquire host-specific virulence genes is intriguing. They can obtain these genes either through horizontal chromosome transfer between strains or through a sequence of loss/gain events or mutations in effector proteins. Therefore, it is conceivable that isolates belonging to a particular formae speciales may share different evolutionary histories leading to their current virulent states (Biju et al., 2016; Henry et al., 2021). This further underlines the complex relationship between genomic diversity and virulence in Foc strains.

As part of our comprehensive investigation to decipher the underlying genetic basis of Foc pathogenicity, we selected two representative strains from each pathogenetic group for long read sequencing: IARI-5175 (virulent) and F-00845 (avirulent). Combining data generated through Nanopore technology and HiC, we were able to achieve scaffold-level ordering of the genomes. The assembled genome sizes were estimated at 46.8 Mb and 42.4 Mb, respectively. We subsequently performed a whole genome syntenic analysis for both Foc strains using the reference genome of *F. oxysporum* f. sp. lycopersici 4287. Interestingly, despite showing a high degree of similarity in the core genomes, we observed very low sequence similarity in the LS region of Fol (Chromosome-3, 6, 14, 15 and parts of chromosome-1 and 2) compared to both strains of Foc (IARI-5175 and F-00845). The comparative genomic analysis between the prototypical virulent and avirulent strains, IARI-5175 and F-00845 respectively, further illustrates the significant genomic variation within the Foc species. This analysis allowed us to partition the genome into two categories: eleven highly conserved core chromosomes, and lineage-specific genomic regions. Notably, in the virulent Foc strain IARI-5175, we identified two additional lineage-specific chromosomes, u000000192 (1.33 Mb) and u000000273 (1.34 Mb). The difference in genome sizes and protein-coding genes between these two strains underscores the genetic complexity within Foc species and its potential link to pathogenicity traits. The alignment with the reference genome of Fol4287 allowed for the assembly of the Foc strains into core chromosomes, revealing the absence of syntenic regions corresponding to accessory chromosomes of Fol4287. The absence of these accessory chromosomes could be a crucial determinant of the observed variability in virulence among the strains.

The discovery of lineage-specific genomic regions in IARI-5175 and F-00845, with their unique genes, highlights the potential role of these regions in influencing strain-specific characteristics, potentially including pathogenicity. The large number of genes unique to IARI-5175 located within its accessory genome, may point towards the presence of specific genetic components linked to its virulence, which are absent in the avirulent F-00845. Such genes could be potential targets for future research focused on combating Fusarium diseases. The presence of a higher proportion of repeat sequences in the IARI-5175 genome suggests a potentially more dynamic genome evolution. Repeat sequences have been known to induce genomic rearrangements, contributing to genome plasticity, a feature often associated with pathogenicity in fungi (Ma et al., 2010). The different distribution and abundance of retroelements and transposons also imply that the two genomes may have undergone different evolutionary paths, potentially driven by different ecological pressures and host interactions.

In conclusion, the findings from this study offer a comprehensive overview of the genetic diversity, evolutionary trajectories, and molecular underpinnings of virulence in *F. oxysporum* f. sp. carthami. Our research underscores the importance of continued exploration into the genetic basis of pathogenicity within this species complex. Such knowledge can support the development of effective strategies to manage and control the spread of these pathogens, ultimately benefiting agricultural production and food security.

Our next major goal was to compare the genomes of virulent and avirulent Foc strains on a broad scale. Plant pathogens carry an assortment of secretory proteins that allow them to invade hosts and challenge their immune systems. Plants, in response, deploy a two-layered defense system. The first layer involves the recognition of pathogen-associated molecular patterns (PAMPs), triggering a defense mechanism called Pathogen Triggered Immunity (PTI) in the host plant. To suppress PTI and continue host invasion, pathogens secrete small, cysteine-rich effector proteins to further the infection process. Fungal genomes have evolved to compartmentalize and accelerate the evolutionary rate of genes encoding these effectors, leading to significant sequence diversity among different formae speciales.

In our investigation, we identified and compared genes coding for effector proteins in virulent (IARI-5175) and avirulent (F-00845) Foc strains. Effector proteins play crucial roles in host-pathogen interactions, often enabling the pathogen to overcome host defenses or modulate host responses. We found a higher number of effectors in the more virulent strain, IARI-5175, indicating a potential correlation between effector repertoire size and virulence. Further, a majority of these effectors were unique to IARI-5175, suggesting that they might be associated with its heightened virulence.

A Gene Ontology (GO) enrichment analysis of these unique effectors pointed to an overrepresentation of genes related to catalytic activity, pectin lysis, and carbohydrate/ derivative binding proteins. Gene Ontology (GO) analysis of these unique effectors pointed to their potential involvement in carbohydrate binding, cell wall degradation, host cell penetration, metabolic processes, and interspecies interaction between organisms, consistent with the roles of fungal effectors in manipulating host cellular processes and defenses (Lo Presti et al., 2015; Sun et al., 2022). Cellular component of gene ontology suggests that at least 10 effectors are located in the extracellular region of the cell. This gives an indication that the effectors are secreted by Foc in the host cell to suppress the plant immunity.

We also examined the expression patterns of these effectors in control mycelia, and in Foc-infected resistant and susceptible safflower accessions. Although we primarily focused on secretory effector proteins with signal peptide sequences, it is worth noting that some apoplastic effector proteins lack these sequences and are secreted via unconventional methods (Sperschneider et al., 2018). Newer effector prediction tools can also detect these unconventionally secreted effectors. As such, we have provided a list of effectors lacking signal peptide sequences for future investigations to potentially uncover additional genes crucial to Foc pathogenicity. Interestingly, we observed considerable variation in the expression of these effectors upon Foc infection in resistant and susceptible safflower roots. The differential expression patterns suggest that the pathogen’s response to host defense is nuanced and can adapt based on the resistance level of the host. This finding strengthens our understanding of the dynamic interplay between the host and pathogen during infection.

We also categorized effectors into three classes based on their expression profiles in resistant and susceptible hosts. Effectors in Class-I, upregulated in susceptible hosts but down regulated in resistant ones, might be crucial for overcoming host defenses. Those in Class-II, with inverse expression patterns, could be essential for the pathogen’s survival in resistant hosts. Class-III effectors, with increased expression in both hosts, might be vital for the basic pathogenicity of the fungus. These classifications could guide future functional studies on these effectors and their roles in fungal pathogenicity. In conclusion, our study unravels the genomic and effector protein differences between virulent and avirulent Foc strains, highlighting the potential roles of repeat sequences and unique effector proteins in virulence. These findings contribute to our understanding of *F. oxysporum* pathogenicity and offer potential targets for disease control strategies. However, further studies are required to functionally validate the roles of these unique effectors in the interaction between *F. oxysporum* and its hosts.

### 4.1 CAZyme repertoire in Foc strains

Much like other fungal pathogens, Foc secretes an array of diverse cell wall degrading enzymes during host invasion. The present study provides a comprehensive analysis of CAZymes distribution in the genomes of two different Fusarium strains - virulent IARI-5175 and avirulent F-00845. We utilized dbCAN to identify the potential CAZyme candidates that could function as CWDEs, a critical component contributing to the virulence of plant pathogens. CAZyme families operate in a coordinated manner to facilitate cell wall modification, biosynthesis, and degradation. Thus, CAZyme classes can also be grouped based on their action on different components of the cell wall. Glycoside Hydrolases (GH) are predominantly involved in cellulose degradation, GH and Carbohydrate Esterases (CE) are crucial for hemicellulose degradation, GH, CE, and Polysaccharide Lyases (PL) participate in pectin degradation, and Auxiliary Activities (AA) are integral for lignin degradation. A notable portion of these enzymes was localized on core chromosomes, suggesting that core chromosomal regions may play a pivotal role in maintaining the functionality and adaptability of these strains. A unique observation was the substantial number of CAZymes unique to IARI-5175 (301), of which 162 GHs, 20 CEs, 21 PLs, 37 AAs, and 9 CBMs were only present in the IARI-5175 strain. This result indicates that the additional repertoire of CAZymes in IARI-5175 could contribute to its enhanced virulence, as some of these enzymes, specifically GHs, are known for their role in the degradation of plant cell wall components, a crucial step in pathogenesis. In contrast, F-00845 had a larger number of strain-specific CAZymes but lacked unique GT class CAZymes, suggesting a potential lack of some pathogenesis-related mechanisms.

To understand the role of these CAZymes in Foc pathogenicity, we identified unique CAZyme coding genes present only in the virulent IARI-5175 strain. A total of 301 unique CAZyme genes were found exclusively in the IARI-5175 strain. Distinct CAZyme families belonging to the GH class, followed by AA, PL, CE, and CBM, were observed in IARI-5175. These CAZymes could potentially contribute to Foc virulence by facilitating successful host invasion. The differential expression analysis revealed that a subset of the unique IARI-5175 CAZymes was differentially regulated during interactions with safflower. This differential expression in a host-dependent manner suggests that these enzymes may have a role in adapting to different host environments and may contribute to pathogenesis. Furthermore, the differential regulation of CAZymes in the resistant disease interaction may imply the activation of alternative metabolic pathways in response to host resistance mechanisms. Previous transcriptome analysis on different races of *F. oxysporum* also highlighted that each race possesses unique sets of CAZymes that exhibit differential expression during wilt disease (Achari et al., 2021).

The findings of our study also shed light on an intriguing aspect of CAZyme regulation during disease interaction. We identified a set of 44 CAZymes that, while unexpressed in the fungal mycelia, were upregulated during the compatible interaction. This suggests their potential role in establishing disease. Interestingly, the expression of three genes showed a negative correlation, suggesting a conditional expression based on the type of interaction. A particularly noteworthy finding was revealed through our comparison of differentially expressed genes (DEGs) in the incompatible interaction with those in the compatible interaction. A gene coding for a multi modular CAZyme having CBM67 and GH78 modules was found to be expressed exclusively in the susceptible host and showed no expression in the resistant host or control Foc mycelial tissue. Rhamnose, a deoxy sugar present in plant cell walls, is a known target of CBM67 and GH78, thus this CAZyme seems to play a pivotal role in degrading cell walls and contributes to the pathogenesis of Foc.

Finally, in our study, we did not find any differentially expressed CAZyme-encoding genes within the accessory genome of the virulent strain during interaction with either susceptible or resistant host plants. This result prompts further investigation to understand if expression of these genes is conditional and whether they play any significant role in pathogenesis. In summary, our study contributes to a better understanding of the role of CAZymes in the pathogenicity of Fusarium strains, providing a foundation for future functional studies. Furthermore, the identified unique CAZymes in the virulent strain could be potential targets for developing disease control strategies. Nevertheless, the study’s limitations are evident, as functional validation of the predicted CAZymes and understanding the mechanistic details of their role in pathogenesis is yet to be done. Additionally, the role of the unexpressed CAZymes in the fungal mycelia needs to be unravelled, potentially in different host-pathogen interaction conditions.

### 4.2 Secondary metabolites

Fungal pathogens produce structurally diverse secondary metabolites that play a crucial role in communication with other coexisting microbes, and virulence factors. Apart from this, they also have significant pharmacological relevance as antimicrobial agents and other bioactive compounds. In our study, we identified secondary metabolites in both Foc strains and compared them to understand the differences between virulent and avirulent strains. A salient finding of this comparative genomic analysis was the discrepancy in the number of secondary metabolite gene clusters between the virulent strain Foc-IARI-5175 and the avirulent strain F-00845. The virulent strain harbored fewer gene clusters, suggesting that the composition, rather than the quantity, of secondary metabolite genes could be a crucial determinant of virulence. This highlights the complex nature of fungal pathogenicity, which seems to be dictated not just by the mere presence of certain genes, but also their organization and interaction within the genomic landscape. Fungal invasion is due to a concerted action of many different factors and a previous study on *Aspergillus fumigatus* also revealed that secondary metabolites are not the only factors responsible for virulence and only a subset of secondary metabolites are known to contribute to fungal pathogenicity (Bignell et al., 2016). It will therefore be important to reveal the secondary metabolites that actually play an active role in Foc’s pathogenesis in safflower by additional functional studies. The distribution of gene clusters was another interesting aspect. In the virulent strain IARI-5175, the gene clusters were exclusively located in the core genome, implying their essential role in the basic survival and functioning of the fungus. On the other hand, the presence of some gene clusters in the accessory genome of the avirulent strain F-00845 might suggest a potential for genomic plasticity and adaptability, which could have implications for the evolution of fungal virulence.

One of the key outcomes was the identification of mycotoxin-producing gene clusters in both strains, with a potential to produce different toxins. Given the known role of mycotoxins as virulence factors in plant-pathogenic fungi, the presence and differential expression of these toxin-producing gene clusters could significantly contribute to the overall pathogenicity and host specificity of the strains. Our analysis revealed that genes coding for fusaric acid were present in both the Foc strains whereas beauvericin and ACT toxin were uniquely identified in IARI-5175 and F-00845 strains, respectively. Fusaric acid (FSA) is shown to be produced by several Fusarium species and is associated with pathogenicity of *F. oxysporum* (Venter & Steyn, 1998). Knockout mutation genes involved in fusaric acid biosynthesis in *F. oxysporum* results in highly reduced virulence (Yang et al., 2023; Phasha et al., 2021).

In our study, we utilized transcriptome data to scrutinize the gene expressions associated with fusaric acid biosynthesis. RNAseq analysis shed light on the differential expressions of secondary metabolite genes during both compatible (Foc interacting with a susceptible host) and incompatible interactions (Foc interacting with a resistant host). This analysis indicates that the fungus modifies its secondary metabolite gene expression in response to the host plant’s resistance or susceptibility, presenting a compelling picture of the complex host-pathogen dynamics at play. We found the expression patterns of genes related to beauveric in and fusaric acid, two mycotoxins, especially noteworthy. We observed differential regulation in 12 out of 22 genes of the fusaric acid biosynthetic cluster during the susceptible interaction, while only three genes exhibited differential expression during the resistant interaction. Notably, gene Foc04.g5197 displayed upregulation in the compatible interaction between Foc and safflower, while its expression was down regulated in the incompatible interaction. This gene demonstrated a high degree of identity with FUB11, a major facilitator superfamily (MFS) transporter of *Fusarium fujikuroi*. Given the cytotoxic nature of fusaric acid (FSA) to fungal cells, fungi have developed mechanisms to reduce its toxicity. One such method relies on FUB11 to transport FSA outside the cell, while another strategy involves converting FSA to a less toxic compound, fusarinolic acid. Thus, the observed down regulation of FUB11 during the incompatible interaction could potentially inhibit the extracellular export of the phytotoxic FSA. This mechanism could be one of the factors contributing to the wilt resistance observed in the Sharda variety.

On the contrary, in the beauvericin biosynthetic gene cluster, we identified two genes showing upregulation and another two that displayed down regulation in the compatible interaction. Among the down regulated genes was one encoding the BEA synthetase, which is reported to be critical for beauvericin synthesis. On the other hand, the upregulated gene was coding for a repressor protein, which inhibits the synthesis of beauvericin. This observed inverse correlation in the expression of specific genes within the beauvericin biosynthetic gene cluster during compatible and incompatible interactions suggests a potential regulatory mechanism for beauvericin biosynthesis dependent on the host’s susceptibility status. This regulation could be crucial for the fungus to fine-tune its virulence and survival tactics, thereby facilitating successful host colonization. Transcriptional reprogramming also hints that the mycotoxin beauvericin may not be necessary in the later stages of Fusarium wilt disease by Foc. Nevertheless, additional investigations are necessary to understand the role of beauvericin in the early stages of Fusarium wilt disease. Previous research on *Fusarium subglutinans* and *Fusarium sacchari* has implicated the potential role of beauvericin in sugarcane pathogenicity (Mohammad et al., 2021).

Moreover, we identified a gene cluster associated with the production of ACT toxin in the avirulent Foc strain F-00845. To date, no reports suggest that *F. oxysporum* employs ACT toxin to enhance its virulence. However, ACT is a known host-selective toxin produced by *Alternaria alternata*, playing a crucial role in its pathogenicity (Huang et al., 2022). A recent investigation on *A. alternata* demonstrated that ACT biosynthesis requires essential genes such as hydrolase, enoyl reductase, HMG CoA hydrolase, polyketide synthase, acyl-CoA synthetase, enoyl-CoA hydratase, and Zn (II)2 cys6 transcription factor (Huang et al., 2022). Upon closer examination, we discovered that the ACT-gene cluster in Foc strain F-00845 contained genes encoding polyketide synthase, aldehyde reductase, alcohol dehydrogenase, oxidoreductase, and methyl transferase. However, it lacked other potential key genes for ACT biosynthesis, such as HMG CoA hydrolase, acyl-CoA synthetase, enoyl-CoA hydratase, and Zn (II)2 cys6 transcription factor. Therefore, it appears that Fusarium may be in the process of acquiring these genes, possibly as part of an adaptive mechanism against the resistance introduced in crop plants. In summary, our study illuminates the multifaceted role of secondary metabolite gene clusters in the pathogenesis of plant-pathogenic fungi. By deciphering the intricate interplay between the fungal genome, secondary metabolite gene clusters, and host plant interactions, it offers potential avenues for developing innovative disease management strategies that target these specific metabolic pathways. This could potentially contribute to more sustainable and efficient agricultural practices in the future.

### 4.3 Transcriptional reprogramming during wilt disease

To further analyze the changes in gene expression associated with Foc virulence, RNA seq data of control fungal mycelia, resistant host infected with Foc (RI) and Susceptible host infected with Foc (SI) were analyzed. The high degree of correlation observed across biological replicates underscores the robustness and reliability of our experimental procedures, thus adding validity to the findings of this study. A total of 5694 and 2210 DEGs were identified in susceptible and resistant hosts, respectively. We observed that a large number of *F. oxysporum* f. sp. ciceris (Foc) genes exhibited differential expression patterns during infection in the susceptible host. This extensive transcriptional reprogramming highlights the complexity of the pathogen-host interaction, where the pathogen manipulates its gene expression to thrive in the host environment. In previous studies, transcriptome analysis has been utilized to describe gene expression changes in susceptible and resistant host in response to invading *F. oxysporum* and led to identification of R genes that mediate resistance in the host (Silvia Sebastiani et al., 2017; Sun et al., 2022). In another transcriptomic study novel effectors of the pathogen were discovered by analysing the differentially expressed genes in *F. oxysporum* f. sp. *medicaginis* (Thatcher et al., 2016).

The GO enrichment analysis provided valuable insights into the functional implications of these differentially expressed genes (DEGs). The up regulated genes in the susceptible host were primarily associated with transport and localization functions. This likely points towards an increased need for the transport of various organic molecules and ions, which could be a critical aspect of the pathogen’s strategy to exploit the host resources and establish successful infection. Down regulated genes in the susceptible host were associated predominantly with small molecule metabolic processes, mitotic cell cycle metabolic processes, and DNA metabolic processes. This could indicate a shift in Foc’s metabolic activity during infection, possibly to divert resources towards other processes that may aid in successful colonization.

Interestingly, in the resistant host, we observed a lower number of DEGs. The nature of up regulated genes was similar to that of the susceptible host, further reinforcing the role of transport-related genes in the pathogen’s infection strategy. However, the down regulated genes in the resistant host included those related to metabolic and transport processes, hinting at a different interaction dynamic. It’s plausible that the resistant host may exert some form of suppression effect on the pathogen, resulting in the down regulation of these genes.

The comparative analysis of DEGs in susceptible and resistant hosts revealed genes with both similar and inverse expression patterns. These genes could potentially represent key elements in the host-pathogen interaction dynamics. The shared up regulated genes, primarily associated with metabolic and catabolic processes, might represent fundamental requirements of the pathogen for host colonization, regardless of the host’s susceptibility status. On the other hand, the inversely correlated genes could underpin the differential responses of the pathogen in susceptible and resistant hosts. It is noteworthy that many of the genes that were up regulated in the susceptible host but down regulated in the resistant host were related to transporter activity. This may suggest that the resistant host somehow interferes with the pathogen’s transport mechanisms, thereby inhibiting its successful colonization.

Overall, these findings underscore the intricate dynamics of the interaction between Foc and its host plants. They highlight potential molecular strategies employed by the pathogen, which could vary significantly depending on the host’s susceptibility. These insights could serve as a foundation for future studies aiming to understand the molecular mechanisms underlying host-pathogen interactions further and could contribute to the development of novel strategies for managing Fusarium wilt in crops.

## Acknowledgments

MA laboratory is funded by Delhi University, Institute of Excellence’s Faculty Research Program Grant for the year 2022 (Sanction No./IoE/2021/12/ 104 FRP). Aabha, and Vijay Laxmi are also thankful to Council of Scientific and Industrial Research (CSIR), India for their research fellowships.

## Authors contributions

MA, KP and A conceived the project. A and VL performed the pathogenicity testing and fungal growth rate measurement. SS and BS performed morphological analysis of fungal strains A constructed the transcriptomic and genomic libraries. KP conducted the genome assembly, transcriptome studies. KP and A performed comparative genomic and transcriptomic analysis. All authors read and approved the final version of the manuscript.

