## Supplementary Material for "Comparative Genomic and Transcriptomic Analysis of Avirulent and Virulent Strains of *Fusarium oxysporum* f. sp. *carthami*: Insights into Pathogenesis and Virulence Determinants in Safflower Infections"

**Supplementary Figure**

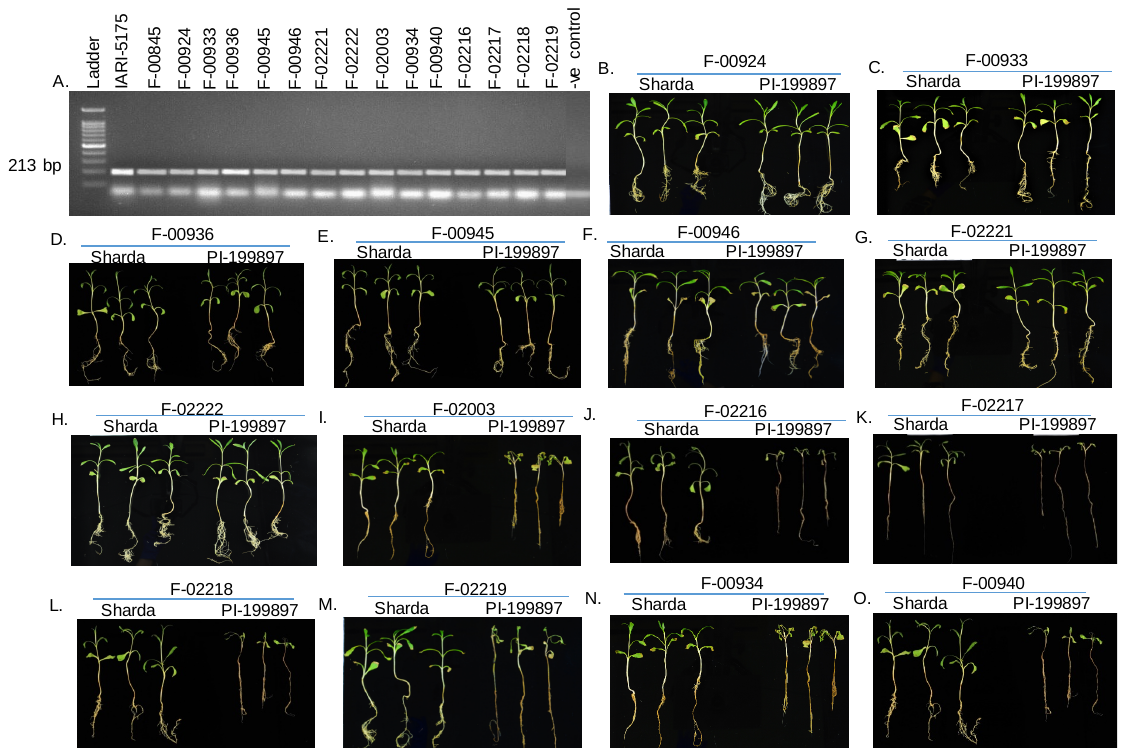

**Supplementary Figure S1:** Confirmation and pathogenicity testing of Foc strains: A) Confirmation of Foc strains with amplification of Foc specific sequence with SCAR marker. B-H) Foc strains showing avirulent phenotype with both Sharda and PI-199897. I-O) Foc strains showing avirulent and virulent phenotype with Sharda and PI-199897 respectively.

**Supplementary Tables**

**Table S1:** List of Foc strains used in the present study

| **S. No.** | **Strain ID** | **S. No.** | **Strain ID** |
| --- | --- | --- | --- |
| 1 | F-00845 | 9 | F-02003 |
| 2 | F-00924 | 10 | F-02216 |
| 3 | F-00933 | 11 | F-02217 |
| 4 | F-00934 | 12 | F-02218 |
| 5 | F-00936 | 13 | F-02219 |
| 6 | F-00940 | 14 | F-02221 |
| 7 | F-00945 | 15 | F-02222 |
| 8 | F-00946 | 16 | IARI-5175 |

**Table S2:** Details of repeat elements present in virulent and avirulent Foc strains.

| **Name of Repeat** | **IARI-5175** | | | **F-00845** | | |
| --- | --- | --- | --- | --- | --- | --- |
|  | **Number of element** | **Length occupied** | **Percentage of  sequence** | **Number of element** | **Length occupied** | **Percentage of  sequence** |
| Retroelements | 829 | 977129 bp | 2.09% | 162 | 145698 bp | 0.34% |
| SINEs: | 0 | 0 bp | 0.00% | 0 | 0 bp | 0.00% |
| Penelope | 0 | 0 bp | 0.00% | 0 | 0 bp | 0.00% |
| LINEs: | 311 | 398600 bp | 0.85% | 45 | 28561 bp | 0.07% |
| CRE/SLACS | 15 | 10296 bp | 0.02% | 0 | 0 bp | 0.00% |
| L2/CR1/Rex | 0 | 0 bp | 0.00% | 0 | 0 bp | 0.00% |
| R1/LOA/Jockey | 0 | 0 bp | 0.00% | 0 | 0 bp | 0.00% |
| R2/R4/NeSL | 0 | 0 bp | 0.00% | 0 | 0 bp | 0.00% |
| RTE/Bov-B | 0 | 0 bp | 0.00% | 0 | 0 bp | 0.00% |
| L1/CIN4 | 0 | 0 bp | 0.00% | 0 | 0 bp | 0.00% |
| LTR elements: | 518 | 578529 bp | 1.24% | 117 | 117137 bp | 0.28% |
| BEL/Pao | 0 | 0 bp | 0.00% | 0 | 0 bp | 0.00% |
| Ty1/Copia | 205 | 317123 bp | 0.68% | 3 | 318 bp | 0.00% |
| Gypsy/DIRS1 | 313 | 261406 bp | 0.56% | 114 | 116819 bp | 0.27% |
| Retroviral | 0 | 0 bp | 0.00% | 0 | 0 bp | 0.00% |
| DNA transposons | 1617 | 1977287 bp | 4.23% | 58 | 15498 bp | 0.04% |
| hobo-Activator | 612 | 853831 bp | 1.83% | 16 | 3707 bp | 0.01% |
| Tc1-IS630-Pogo | 596 | 684937 bp | 1.47% | 27 | 9737 bp | 0.02% |
| En-Spm | 0 | 0 bp | 0.00% | 0 | 0 bp | 0.00% |
| MuDR-IS905 | 0 | 0 bp | 0.00% | 0 | 0 bp | 0.00% |
| PiggyBac | 89 | 78219 bp | 0.17% | 8 | 1256 bp | 0.00% |
| Tourist/Harbinger | 0 | 0 bp | 0.00% | 0 | 0 bp | 0.00% |
| Other (Mirage, | 0 | 0 bp | 0.00% | 0 | 0 bp | 0.00% |
| P-element, Trans | ib) |  |  | ib) |  |  |
| Rolling-circles | 56 | 139207 bp | 0.30% | 7 | 3225 bp | 0.01% |
| Unclassified: | 5598 | 2292495 bp | 4.91% | 3043 | 777576 bp | 1.83% |
| Total interspersed repeats |  | 5246911 bp | 11.23% |  | 938772 bp | 2.21% |
| Small RNA: | 12 | 6439 bp | 0.01% | 15 | 1399 bp | 0.00% |
| Satellites: | 0 | 0 bp | 0.00% | 0 | 0 bp | 0.00% |
| Simple repeats | 5210 | 213920 bp | 0.46% | 7517 | 297194 bp | 0.70% |
| Low complexity | 581 | 28988 bp | 0.06% | 745 | 35203 bp | 0.08% |

**Table S3:** **Chromosome wise distribution of CAZymes present in the virulent (V) and avirulent (AV) Foc strains.** The number of CAZymes on core chromosomes (Foc01-Foc11), lineage specific chromosomes of the virulent strain (u000000192 and u000000273) and lineage specific region (LS) of the avirulent strain is listed.

| **Chromosome ID** | **(AAs)** | | **(CBMs)** | | **(CEs)** | | **(GHs)** | | **(GTs)** | | **(PLs)** | |
| --- | --- | --- | --- | --- | --- | --- | --- | --- | --- | --- | --- | --- |
|  | **V** | **AV** | **V** | **AV** | **V** | **AV** | **V** | **AV** | **V** | **AV** | **V** | **AV** |
| Foc01 | 8 | 9 | 2 | 0 | 5 | 3 | 25 | 28 | 9 | 14 | 2 | 1 |
| Foc02 | 7 | 9 | 0 | 0 | 2 | 3 | 17 | 18 | 9 | 12 | 4 | 3 |
| Foc03 | 16 | 17 | 1 | 0 | .1 | 4 | 27 | 27 | 7 | 13 | 1 | 2 |
| Foc04 | 6 | 8 | 2 | 1 | 3 | 3 | 24 | 24 | 8 | 8 | 1 | 0 |
| Foc05 | 3 | 7 | 1 | 1 | 3 | 4 | 17 | 21 | 7 | 9 | 2 | 0 |
| Foc06 | 6 | 7 | 0 | 0 | 1 | 1 | 31 | 34 | 6 | 7 | 1 | 0 |
| Foc07 | 4 | 6 | 1 | 1 | 3 | 2 | 17 | 10 | 8 | 10 | 0 | 0 |
| Foc08 | 3 | 6 | 0 | 0 | 1 | 2 | 7 | 8 | 3 | 8 | 2 | 1 |
| Foc09 | 9 | 14 | 2 | 1 | 4 | 3 | 29 | 31 | 3 | 4 | 1 | 1 |
| Foc10 | 13 | 4 | 1 | 0 | 6 | 4 | 28 | 18 | 1 | 2 | 6 | 1 |
| Foc11 | 10 | 11 | 1 | 0 | 2 | 2 | 17 | 16 | 3 | 3 | 3 | 3 |
| u000000192 | 1 | - | 0 | - | 0 | - | 3 | - | 0 | - | 0 | - |
| u000000273 | 0 | - | 0 | - | 0 | - | 1 | - | 0 | - | 0 | - |
| LS | 2 | 7 | 0 | 0 | 0 | 2 | 7 | 11 | 0 | 4 | 0 | 0 |

**Table S4**: List of genes coding for putative effectors in virulent and avirulent strain of Foc.

| A. **List of genes coding for putative effector proteins in virulent Foc strain** | | | |
| --- | --- | --- | --- |
| Foc01.g1175.t1 | Foc03.g4551.t1 | Foc05.g7754.t1 | Foc09.g10960.t |
| Foc01.g1211.t1 | Foc03.g4641.t1 | Foc05.g7791.t1 | Foc09.g10968.t |
| Foc01.g1557.t1 | Foc03.g4646.t1 | Foc05.g7847.t1 | Foc09.g10970.t |
| Foc01.g1608.t1 | Foc03.g4922.t1 | Foc05.g7950.t1 | Foc09.g10979.t |
| Foc01.g1736.t1 | Foc03.g4947.t1 | Foc05.g7953.t1 | Foc09.g11030.t |
| Foc01.g1760.t1 | Foc03.g4990.t1 | Foc05.g7959.t1 | Foc09.g11037.t |
| Foc01.g1802.t1 | Foc03.g4999.t1 | Foc05.g8031.t1 | Foc09.g11062.t |
| Foc01.g187.t1 | Foc03.g5000.t1 | Foc05.g8054.t1 | Foc09.g11064.t |
| Foc01.g38.t1 | Foc03.g5001.t1 | Foc05.g8057.t1 | Foc09.g11088.t |
| Foc01.g383.t1 | Foc03.g5019.t1 | Foc05.g8062.t1 | Foc09.g11101.t |
| Foc01.g485.t1 | Foc03.g5021.t1 | Foc05.g8102.t1 | Foc09.g11124.t |
| Foc01.g507.t1 | Foc03.g5065.t1 | Foc05.g8133.t1 | Foc09.g11132.t |
| Foc01.g6.t1 | Foc03.g5081.t1 | Foc06.g8201.t1 | Foc09.g11153.t |
| Foc01.g610.t1 | Foc03.g5104.t1 | Foc06.g8234.t1 | Foc09.g11178.t |
| Foc01.g752.t1 | Foc04.g5192.t1 | Foc06.g8258.t1 | Foc09.g11264.t |
| Foc01.g917.t1 | Foc04.g5229.t1 | Foc06.g8270.t1 | Foc09.g11346.t |
| Foc01.g928.t1 | Foc04.g5235.t1 | Foc06.g8273.t1 | Foc09.g11349.t |
| Foc01.g945.t1 | Foc04.g5249.t1 | Foc06.g8275.t1 | Foc09.g11417.t |
| Foc02.g1849.t1 | Foc04.g5283.t1 | Foc06.g8300.t1 | Foc09.g11424.t |
| Foc02.g1851.t1 | Foc04.g5292.t1 | Foc06.g8356.t1 | Foc09.g11471.t |
| Foc02.g1861.t1 | Foc04.g5299.t1 | Foc06.g8359.t1 | Foc09.g11472.t |
| Foc02.g1920.t1 | Foc04.g5548.t1 | Foc06.g8369.t1 | Foc09.g11523.t |
| Foc02.g1954.t1 | Foc04.g5570.t1 | Foc06.g8375.t1 | Foc09.g11540.t |
| Foc02.g1968.t1 | Foc04.g5672.t1 | Foc06.g8725.t1 | Foc09.g11560.t |
| Foc02.g1984.t1 | Foc04.g6070.t1 | Foc06.g8895.t1 | Foc09.g11570.t |
| Foc02.g1990.t1 | Foc04.g6287.t1 | Foc06.g9085.t1 | Foc09.g11588.t |
| Foc02.g2014.t1 | Foc04.g6291.t1 | Foc06.g9187.t1 | Foc09.g11632.t |
| Foc02.g2025.t1 | Foc04.g6355.t1 | Foc06.g9342.t1 | Foc10.g11736.t |
| Foc02.g2026.t1 | Foc04.g6447.t1 | Foc07.g10053.t | Foc10.g11747.t |
| Foc02.g2087.t1 | Foc04.g6511.t1 | Foc07.g10080.t | Foc10.g11759.t |
| Foc02.g2098.t1 | Foc04.g6584.t1 | Foc07.g10191.t | Foc10.g11772.t |
| Foc02.g2104.t1 | Foc04.g6608.t1 | Foc07.g10273.t | Foc10.g11779.t |
| Foc02.g2117.t1 | Foc04.g6611.t1 | Foc07.g10295.t | Foc10.g11781.t |
| Foc02.g2169.t1 | Foc04.g6641.t1 | Foc07.g10297.t | Foc10.g11790.t |
| Foc02.g2313.t1 | Foc04.g6683.t1 | Foc07.g10319.t | Foc10.g11798.t |
| Foc02.g2487.t1 | Foc04.g6718.t1 | Foc07.g10349.t | Foc10.g11847.t |
| Foc02.g2506.t1 | Foc04.g6749.t1 | Foc07.g10365.t | Foc11.g11882.t |
| Foc02.g2740.t1 | Foc05.g6759.t1 | Foc07.g9392.t1 | Foc11.g11926.t |
| Foc02.g2863.t1 | Foc05.g6806.t1 | Foc07.g9532.t1 | Foc11.g11958.t |
| Foc02.g2890.t1 | Foc05.g6851.t1 | Foc07.g9663.t1 | Foc11.g12003.t |
| Foc02.g2962.t1 | Foc05.g6892.t1 | Foc07.g9725.t1 | Foc11.g12010.t |
| Foc02.g2992.t1 | Foc05.g6949.t1 | Foc08.g10393.t | Foc11.g12045.t |
| Foc02.g3010.t1 | Foc05.g6983.t1 | Foc08.g10395.t | Foc11.g12094.t |
| Foc02.g3150.t1 | Foc05.g7036.t1 | Foc08.g10398.t | Foc11.g12125.t |
| Foc02.g3246.t1 | Foc05.g7055.t1 | Foc08.g10446.t | u000000192.g13417 |
| Foc03.g3348.t1 | Foc05.g7169.t1 | Foc08.g10496.t | u000000193.g13513 |
| Foc03.g3494.t1 | Foc05.g7190.t1 | Foc08.g10502.t | u000000194.g13557 |
| Foc03.g3515.t1 | Foc05.g7348.t1 | Foc08.g10536.t | u000000194.g13574 |
| Foc03.g3664.t1 | Foc05.g7432.t1 | Foc08.g10577.t | u000000216.g13781 |
| Foc03.g3742.t1 | Foc05.g7493.t1 | Foc08.g10705.t | u000000217.g13782 |
| Foc03.g3756.t1 | Foc05.g7559.t1 | Foc08.g10775.t | u000000219.g13813 |
| Foc03.g3866.t1 | Foc05.g7589.t1 | Foc08.g10818.t | u000000225.g13848 |
| Foc03.g4157.t1 | Foc05.g7714.t1 | Foc08.g10824.t | u000000226.g13851 |
| Foc03.g4200.t1 | Foc05.g7715.t1 | Foc08.g10876.t | u000000238.g13905 |
| Foc03.g4278.t1 | Foc05.g7716.t1 | Foc08.g10886.t | u000000273.g14028 |
| Foc03.g4515.t1 | Foc05.g7717.t1 | Foc09.g10913.t |  |
| Foc03.g4526.t1 | Foc05.g7734.t1 | Foc09.g10954.t |  |
| B. **List of genes coding for putative effector proteins in avirulent Foc strain** | | | |
| Foc1.g176.t1 | Foc2.g1680.t1 | Foc4.g5219.t1 | Foc8.g8698.t1 |
| Foc1.g223.t1 | Foc2.g1743.t1 | Foc4.g5302.t1 | Foc9.g8971.t1 |
| Foc1.g33.t1_ | Foc2.g1757.t1 | Foc4.g5307.t1 | Foc9.g8980.t1 |
| Foc1.g362.t1 | Foc2.g1764.t1 | Foc5.g5415.t1 | Foc9.g9001.t1 |
| Foc1.g406.t1 | Foc2.g1873.t1 | Foc5.g5658.t1 | Foc9.g9003.t1 |
| Foc1.g455.t1 | Foc2.g2190.t1 | Foc5.g5741.t1 | Foc9.g9131.t1 |
| Foc1.g761.t1 | Foc2.g2448.t1 | Foc5.g5885.t1 | Foc9.g9148.t1 |
| Foc1.g795.t1 | Foc2.g2593.t1 | Foc5.g6030.t1 | Foc9.g9152.t1 |
| Foc1.g812.t1 | Foc3.g2802.t1 | Foc5.g6190.t1 | Foc9.g9173.t1 |
| Foc1.g818.t1 | Foc3.g2805.t1 | Foc5.g6323.t1 | Foc9.g9184.t1 |
| Foc10.g9749.t | Foc3.g2835.t1 | Foc5.g6359.t1 | Foc9.g9238.t1 |
| Foc10.g9784.t | Foc3.g2842.t1 | Foc5.g6361.t1 | Foc9.g9243.t1 |
| Foc10.g9817.t | Foc3.g2879.t1 | Foc5.g6368.t1 | Foc9.g9309.t1 |
| Foc10.g9864.t | Foc3.g3397.t1 | Foc5.g6422.t1 | Foc9.g9401.t1 |
| Foc10.g9874.t | Foc3.g3442.t1 | Foc5.g6426.t1 | Foc9.g9411.t1 |
| Foc11.g10037 | Foc3.g3515.t1 | Foc6.g6475.t1 | Foc9.g9458.t1 |
| Foc11.g10130 | Foc3.g3571.t1 | Foc6.g6481.t1 | Foc9.g9521.t1 |
| Foc11.g10200 | Foc3.g3708.t1 | Foc6.g6553.t1 | Foc9.g9530.t1 |
| Foc11.g10202 | Foc3.g3918.t1 | Foc6.g6570.t1 | tig00000021.g10528.t1 |
| Foc11.g10289 | Foc3.g4009.t1 | Foc6.g6734.t1 | tig00000021.g10583.t1 |
| Foc11.g10343 | Foc3.g4103.t1 | Foc6.g7292.t1 | tig00000021.g10657.t1 |
| Foc11.g10352 | Foc3.g4125.t1 | Foc6.g7444.t1 |  |
| Foc11.g10384 | Foc4.g4190.t1 | Foc7.g7972.t1 |  |
| Foc11.g9995.t | Foc4.g4264.t1 | Foc7.g8187.t1 |  |
| Foc2.g1536.t1 | Foc4.g4507.t1 | Foc7.g8287.t1 |  |
| Foc2.g1614.t1 | Foc4.g4530.t1 | Foc8.g8406.t1 |  |
|  | Foc4.g5187.t1 | Foc8.g8416.t1 |  |

Table S5. Gene ontology of effectors unique to Foc strain IARI-5175

| **Biological processes** | |
| --- | --- |
| Pathways | Number of genes |
| Small molecule metabolic process | 2 |
| Monosaccharide metabolic process | 1 |
| Organic acid metabolic process | 1 |
| Hexose metabolic process | 1 |
| Carboxylic acid metabolic process | 1 |
| Oxoacid metabolic process | 1 |
| Organic substance metabolic process | 2 |
| Metabolic process | 2 |
| Carbohydrate metabolic process | 1 |
| Cellular metabolic process | 1 |
| Pathogenesis | 1 |
| Cellular process | 1 |
| Interspecies interaction between organisms | 1 |
| **Cellular components** | |
| Extracellular region | 10 |
| Cellular anatomical entity | 10 |
| Plasma membrane | 2 |
| Anchored component of membrane | 2 |
| Cell periphery Membrane | 2 |
| Intrinsic component of membrane | 2 |
| Cell wall | 1 |
| External encapsulating structure | 1 |
| **Molecular pathways** | |
| Binding | 3 |
| Catalytic activity | 3 |
| Hydrolase activity, acting on carbon-nitrogen bonds | 1 |
| Lyase activity | 1 |
| Isomerase activity | 1 |
| Carbohydrate binding | 1 |
| Chitin binding | 1 |
| Carbohydrate derivative binding | 1 |
| Cation binding | 1 |
| Metal ion binding | 1 |
| Ion binding | 1 |
| Hydrolase activity | 1 |

Table S6: Gene ontology terms for genes differentially expressed in compatible and incompatible interaction.

| A. Upregulated genes in susceptible infected plants vs control mycelia | |
| --- | --- |
| Biological processes | |
| Pathways | Number of genes |
| Amino acid transmembrane transport | 10 |
| Amino acid transport | 11 |
| Carboxylic acid transmembrane transport | 15 |
| Carboxylic acid transport | 17 |
| Organic acid transport | 17 |
| Anion transmembrane transport | 19 |
| Anion transport | 23 |
| Organic anion transport | 19 |
| Monovalent inorganic cation transport | 15 |
| Transmembrane transport | 50 |
| Ion transmembrane transport | 32 |
| Cation transmembrane transport | 22 |
| Cation transport | 26 |
| Ion transport | 40 |
| Transport | 83 |
| Establishment of localization | 83 |
| Organic substance transport | 56 |
| Localization | 88 |
| Nitrogen compound transport | 45 |
| **Cellular components** | |
| Pathways | Number of genes |
| Intrinsic component of vacuolar membrane | 4 |
| Eisosome | 3 |
| Integral component of vacuolar membrane | 3 |
| Vacuole-mitochondrion membrane contact site | 3 |
| Integral component of plasma membrane | 15 |
| Intrinsic component of plasma membrane | 16 |
| Fungal-type vacuole membrane | 24 |
| Lytic vacuole membrane | 24 |
| Vacuolar membrane | 28 |
| Storage vacuole | 45 |
| Fungal-type vacuole | 45 |
| Lytic vacuole | 45 |
| Vacuole | 47 |
| Plasma membrane | 49 |
| Cell periphery | 65 |
| Bounding membrane of organelle | 44 |
| Whole membrane | 36 |
| Integral component of membrane | 87 |
| Organelle membrane | 54 |
| Molecular functions | |
| Pathways | Number of genes |
| Alcohol transmembrane transporter activity | 3 |
| Potassium ion transmembrane transporter activity | 5 |
| Symporter activity | 11 |
| Solute:cation symporter activity | 9 |
| Amino acid transmembrane transporter activity | 10 |
| Solute:proton symporter activity | 8 |
| Secondary active transmembrane transporter activity | 19 |
| Carboxylic acid transmembrane transporter activity | 14 |
| Organic anion transmembrane transporter activity | 15 |
| Anion transmembrane transporter activity | 18 |
| Active transmembrane transporter activity | 23 |
| Active ion transmembrane transporter activity | 13 |
| Monovalent inorganic cation transmembrane transporter activity | 15 |
| Transmembrane transporter activity | 44 |
| Inorganic molecular entity transmembrane transporter activity | 31 |
| Transporter activity | 47 |
| Cation transmembrane transporter activity | 22 |
| Ion transmembrane transporter activity | 30 |
| Inorganic cation transmembrane transporter activity | 17 |
| B. **Downregulated genes in susceptible infected plants vs control mycelia** | |
| Biological processes | |
| Pathways | Number of genes |
| DNA unwinding involved in DNA replication | 5 |
| Pre-replicative complex assembly | 6 |
| Pre-replicative complex assembly involved in nuclear cell cycle DNA replication | 6 |
| DNA replication initiation | 13 |
| Cell cycle DNA replication initiation | 12 |
| DNA duplex unwinding | 15 |
| DNA geometric change | 15 |
| Mitotic DNA replication | 24 |
| Nuclear DNA replication | 25 |
| Cell cycle DNA replication | 25 |
| DNA repair | 41 |
| Mitotic cell cycle proc. | 75 |
| Carboxylic acid metabolic proc. | 52 |
| Oxoacid metabolic proc. | 53 |
| DNA metabolic proc. | 54 |
| Small molecule metabolic proc. | 87 |
| **Cellular components** | |
| Pathways | Number of genes |
| Ski complex | 3 |
| Eukaryotic translation initiation factor 2B complex | 5 |
| MCM complex | 5 |
| Arp2/3 protein complex | 4 |
| Nuclear pre-replicative complex | 7 |
| Pre-replicative complex | 7 |
| DNA replication preinitiation complex | 8 |
| Replisome | 7 |
| Nuclear replisome | 7 |
| Proteasome regulatory particle | 7 |
| Proteasome accessory complex | 7 |
| Nuclear replication fork | 19 |
| Replication fork | 19 |
| Protein-DNA complex | 19 |
| Actin cortical patch | 14 |
| Endocytic patch | 14 |
| Cortical actin cytoskeleton | 21 |
| Cortical cytoskeleton | 23 |
| Nuclear chromosome | 28 |
| Cytoskeleton | 57 |
| Molecular functions | |
| Pathway | Number of Genes |
| Single-stranded DNA binding | 18 |
| ATP-dependent activity, acting on DNA | 22 |
| Cytoskeletal protein binding | 26 |
| Helicase activity | 25 |
| Catalytic activity, acting on DNA | 32 |
| ATP-dependent activity | 54 |
| Anion binding | 129 |
| ATP binding | 89 |
| Adenyl nucleotide binding | 90 |
| Adenyl ribonucleotide binding | 89 |
| Nucleotide binding | 122 |
| Nucleoside phosphate binding | 122 |
| Small molecule binding | 130 |
| Carbohydrate derivative binding | 106 |
| Ribonucleotide binding | 105 |
| Purine nucleotide binding | 102 |
| Purine ribonucleoside triphosphate binding | 100 |
| Purine ribonucleotide binding | 100 |
| Ion binding | 195 |
| Protein binding | 185 |
| C. **Upregulated genes in resistant infected plants vs control mycelia** | |
| Biological processes | |
| Pathways | Number of genes |
| Transport | 32 |
| Establishment of localization | 32 |
| Transmembrane transport | 24 |
| Organic substance transport | 23 |
| Nitrogen compound transport | 22 |
| Ion transmembrane transport | 18 |
| Anion transmembrane transport | 13 |
| Organic anion transport | 11 |
| Cation transmembrane transport | 11 |
| Organic acid transmembrane transport | 9 |
| Carboxylic acid transport | 9 |
| Import into cell | 6 |
| Amino acid transmembrane transport | 5 |
| Response to ketone | 3 |
| Cellular response to ketone | 3 |
| Monocarboxylic acid transport | 3 |
| L-amino acid transport | 3 |
| L-alpha-amino acid transmembrane transport | 3 |
| Dipeptide transmembrane transport | 2 |
| **Cellular components** | |
| Pathways | Number of genes |
| Integral component of membrane | 31 |
| Intrinsic component of membrane | 31 |
| Integral component of plasma membrane | 7 |
| Plasma membrane region | 5 |
| Plasma membrane of cell tip | 4 |
| Molecular functions | |
| Pathways | Number of genes |
| Transmembrane transporter activity | 22 |
| Transporter activity | 22 |
| Ion transmembrane transporter activity | 16 |
| Inorganic molecular entity transmembrane transporter activity | 14 |
| Anion transmembrane transporter activity | 11 |
| Organic anion transmembrane transporter activity | 10 |
| Organic acid transmembrane transporter activity | 9 |
| Cation transmembrane transporter activity | 9 |
| Amide transmembrane transporter activity | 5 |
| Amino acid transmembrane transporter activity | 5 |
| Secondary active transmembrane transporter activity | 5 |
| L-amino acid transmembrane transporter activity | 3 |
| Dipeptide transmembrane transporter activity | 2 |
| Vitamin transmembrane transporter activity | 2 |
| D. **Downregulated genes in resistant infected plants vs control mycelia** | |
| Biological processes | |
| Pathways | Number of genes |
| Fructose 2,6-bisphosphate metabolic process | 2 |
| Vitamin transport | 3 |
| Iron coordination entity transport | 3 |
| Monocarboxylic acid catabolic process | 5 |
| Carboxylic acid catabolic process | 6 |
| Small molecule catabolic process | 9 |
| Anion transmembrane transport | 8 |
| Monocarboxylic acid metabolic process | 8 |
| Cellular carbohydrate metabolic process | 7 |
| Ion transmembrane transport | 13 |
| Carbohydrate metabolic process | 10 |
| Transmembrane transport | 18 |
| Ion transport | 15 |
| Oxoacid metabolic process | 15 |
| Organic acid metabolic process | 15 |
| Carboxylic acid metabolic process | 14 |
| Small molecule metabolic process | 22 |
| Cellular components | |
| Pathways | Number of genes |
| Integral component of fungal-type vacuolar membrane | 2 |
| Storage vacuole | 23 |
| Fungal-type vacuole | 23 |
| Fungal-type vacuole membrane | 11 |
| Lytic vacuole | 23 |
| Lytic vacuole membrane | 11 |
| Vacuole | 23 |
| Vacuolar membrane | 12 |
| Plasma membrane | 22 |
| Cell periphery | 26 |
| Whole membrane | 13 |
| Integral component of membrane | 29 |
| Molecular pathways | |
| Pathways | Number of genes |
| 6-phosphofructo-2-kinase activity | 2 |
| Phosphofructokinase activity | 2 |
| Modified amino acid transmembrane transporter activity | 2 |
| Carbohydrate phosphatase activity | 2 |
| Ferric-chelate reductase (nadph) activity | 2 |
| Vitamin transmembrane transporter activity | 2 |
| Oxidoreductase activity, oxidizing metal ions, nad or nadp as acceptor | 2 |
| Nucleobase transmembrane transporter activity | 2 |
| Hydrolase activity, hydrolyzing o-glycosyl compounds | 5 |
| Glucosidase activity | 3 |
| Hydrolase activity, acting on glycosyl bonds | 5 |
| Organic anion transmembrane transporter activity | 7 |
| Carboxylic acid transmembrane transporter activity | 5 |
| Anion transmembrane transporter activity | 7 |
| Transmembrane transporter activity | 15 |
| Ion transmembrane transporter activity | 11 |
| Transporter activity | 15 |
| Oxidoreductase activity | 12 |
| E. **DEGs upregulated in both susceptible and resistant interaction** | |
| Biological Processes | |
| Pathways | Number of genes |
| Galacturonan metabolic process | 7 |
| Pectin metabolic process | 7 |
| Pectin catabolic process | 7 |
| Polysaccharide catabolic process | 9 |
| Polysaccharide metabolic process | 9 |
| Carbohydrate catabolic process | 9 |
| Carbohydrate metabolic process | 10 |
| External encapsulating structure organization | 7 |
| Cell wall organization or biogenesis | 8 |
| Cell wall organization | 7 |
| Macromolecule catabolic process | 9 |
| Organic substance catabolic process | 10 |
| Catabolic process | 10 |
| Cellular component organization | 10 |
| Cellular component organization or biogenesis | 10 |
| DNA packaging | 2 |
| Sterigmatocystin metabolic process | 2 |
| Sterigmatocystin biosynthetic process | 2 |
| Organic heteropentacyclic compound metabolic process | 2 |
| Organic heteropentacyclic compound biosynthetic process | 2 |
| Acetaldehyde catabolic process | 1 |
| Acetaldehyde metabolic process | 1 |
| Toxin biosynthetic process | 2 |
| Toxin metabolic process | 2 |
| Chromatin remodeling | 2 |
| Aspartate family amino acid metabolic process | 2 |
| Secondary metabolite biosynthetic process | 3 |
| DNA conformation change | 2 |
| de novo L-methionine biosynthetic process | 1 |
| Chromosome organization | 3 |
| Secondary metabolic process | 3 |
| Sporocarp development involved in asexual reproduction | 1 |
| Sporulation resulting in formation of a cellular spore | 2 |
| Arabinan metabolic process | 1 |
| Arabinan catabolic process | 1 |
| Primary alcohol catabolic process | 1 |
| Nitrate metabolic process | 1 |
| Nitrate assimilation | 1 |
| Anatomical structure formation involved in morphogenesis | 2 |
| Ethanol catabolic process | 1 |
| Chromatin organization | 2 |
| Threonine catabolic process | 1 |
| Mitotic chromosome condensation | 1 |
| Cellular components | Cellular components |
| Pathways | Number of genes |
| Extracellular region | 8 |
| Molecular Functions | |
| Pathways | Number of genes |
| Carbon-oxygen lyase activity, acting on polysaccharides | 6 |
| Carbon-oxygen lyase activity | 6 |
| Lyase activity | 7 |
| Pectate lyase activity | 3 |
| Pectin lyase activity | 2 |
| Chitin deacetylase activity | 1 |
| Cysteine synthase activity | 1 |
| Glutamate-ammonia ligase activity | 1 |
| Hydrolase activity, hydrolyzing O-glycosyl compounds | 3 |
| Acetylesterase activity | 1 |
| Ammonia ligase activity | 1 |
| Hydrolase activity, acting on glycosyl bonds | 3 |
| Acid-ammonia (or amide) ligase activity | 1 |
| Pectinesterase activity | 1 |
| Short-chain carboxylesterase activity | 1 |
| Arabinan endo-1,5-alpha-L-arabinosidase activity | 1 |
| Polygalacturonase activity | 1 |
| Oxo-acid-lyase activity | 1 |
| Aldehyde-lyase activity | 1 |
| F. **DEGs downregulated in both susceptible and resistant interaction** | |
| Biological Processes | |
| Pathways | Number of genes |
| Membrane docking | 5 |
| Oligopeptide transmembrane transport | 3 |
| Endoplasmic reticulum localization | 3 |
| Endoplasmic reticulum-plasma membrane tethering | 3 |
| Organelle localization by membrane tethering | 5 |
| Oligopeptide transport | 3 |
| Peptide transport | 3 |
| Glutathione transport | 2 |
| Glutathione transmembrane transport | 2 |
| Tripeptide transmembrane transport | 2 |
| Tripeptide transport | 2 |
| Response to inorganic substance | 4 |
| Molecular Functions | |
| Pathways | Number of genes |
| Lipid binding | 7 |
| Protein-membrane adaptor activity | 4 |
| Peptide transmembrane transporter activity | 3 |
| Phospholipid binding | 5 |
| Amide transmembrane transporter activity | 3 |
| G. **DEGs upregulated in susceptible interaction and downregulated in resistant interaction** | |
| Biological Processes | |
| Pathways | Number of genes |
| Cation transport | 10 |
| Transmembrane transport | 13 |
| Ion transport | 12 |
| Transport | 21 |
| Monovalent inorganic cation transport | 7 |
| Establishment of localization | 21 |
| Localization | 22 |
| Inorganic cation transmembrane transport | 7 |
| Inorganic ion transmembrane transport | 7 |
| Metal ion transport | 5 |
| Cation transmembrane transport | 7 |
| Ion transmembrane transport | 8 |
| Proton transmembrane transport | 5 |
| Purine-containing compound transmembrane transport | 2 |
| Copper ion import | 2 |
| Cellular Components | |
| Pathways | Number of genes |
| Plasma membrane | 15 |
| Storage vacuole | 13 |
| Lytic vacuole | 13 |
| Fungal-type vacuole | 13 |
| Vacuole | 14 |
| Cell periphery | 17 |
| Vacuolar membrane | 10 |
| Intrinsic component of membrane | 23 |
| Integral component of membrane | 22 |
| Fungal-type vacuole membrane | 8 |
| Lytic vacuole membrane | 8 |
| Bounding membrane of organelle | 13 |
| Whole membrane | 12 |
| Intrinsic component of plasma membrane | 5 |
| Integral component of plasma membrane | 4 |
| Organelle membrane | 15 |
| Molecular Functions | |
| Pathways | Number of genes |
| Monovalent inorganic cation transmembrane transporter activity | 7 |
| Transporter activity | 11 |
| Secondary active transmembrane transporter activity | 6 |
| Active ion transmembrane transporter activity | 6 |
| Active transmembrane transporter activity | 7 |
| Inorganic cation transmembrane transporter activity | 7 |
| Transmembrane transporter activity | 10 |
| Solute:proton symporter activity | 4 |
| Solute:cation symporter activity | 4 |
| Cation transmembrane transporter activity | 7 |
| Inorganic molecular entity transmembrane transporter activity | 8 |
| Symporter activity | 4 |
| Ion transmembrane transporter activity | 8 |
| Proton transmembrane transporter activity | 5 |
| Ferric-chelate reductase (nadph) activity | 2 |
| Ferric-chelate reductase activity | 2 |
| Oxidoreductase activity, oxidizing metal ions, nad or nadp as acceptor | 2 |
| Potassium ion transmembrane transporter activity | 2 |
| Oxidoreductase activity, oxidizing metal ions | 2 |
| Metal ion transmembrane transporter activity | 3 |
| Ion binding | 18 |
| Amide transmembrane transporter activity | 2 |
| Carbohydrate:proton symporter activity | 2 |
| Glucose transmembrane transporter activity | 2 |
| Carbohydrate:cation symporter activity | 2 |
| Monosaccharide transmembrane transporter activity | 2 |
| Hexose transmembrane transporter activity | 2 |
| Cyclin-dependent protein serine/threonine kinase regulator activity | 2 |
| Sugar transmembrane transporter activity | 2 |
| H. **DEGs downregulated in susceptible interaction and upregulated in resistant interaction** | |
| Biological Processes | |
| Pathways | Number of genes |
| Carbohydrate metabolic process | 8 |
| Small molecule metabolic process | 14 |
| Transmembrane transport | 11 |
| Cellular Components | |
| Pathways | Number of genes |
| Storage vacuole | 16 |
| Lytic vacuole | 16 |
| Fungal-type vacuole | 16 |
| Vacuole | 16 |
| Fungal-type vacuole membrane | 8 |
| Lytic vacuole membrane | 8 |
| Plasma membrane | 12 |
| Cell periphery | 15 |
| Vacuolar membrane | 8 |
| Integral component of fungal-type vacuolar membrane | 2 |
| Intrinsic component of fungal-type vacuolar membrane | 2 |
| Integral component of vacuolar membrane | 2 |
| Intrinsic component of vacuolar membrane | 2 |
| Extracellular region | 4 |
| Intrinsic component of membrane | 17 |
| Molecular Functions | |
| Pathways | Number of genes |
| Hydrolase activity, hydrolyzing o-glycosyl compounds | 4 |
| Transporter activity | 9 |
| Anion transmembrane transporter activity | 5 |
| Organic anion transmembrane transporter activity | 5 |
| Ion transmembrane transporter activity | 8 |
| Glucosidase activity | 3 |
| Hydrolase activity | 15 |
| Hydrolase activity, acting on glycosyl bonds | 4 |
| Carbohydrate phosphatase activity | 2 |
| Transmembrane transporter activity | 9 |
| Metalloaminopeptidase activity | 2 |
| Organic hydroxy compound transmembrane transporter activity | 2 |
| Metalloexopeptidase activity | 2 |
| Aminopeptidase activity | 2 |
